# Multi-platform framework for mapping somatic retrotransposition in human tissues

**DOI:** 10.1101/2025.10.07.680917

**Authors:** Seunghyun Wang, Mingyun Bae, Jinhao Wang, Boxun Zhao, Khue Nguyen, Shayna Mallett, Jessica A. Switzenberg, Steven J. Losh, Corinne E. Sexton, Benpeng Miao, Shihua Dong, Xi Zeng, Ziying Wang, Torrin L. McDonald, Camille Mumm, Rohini K. Gadde, Arnaz Maryam Tariq, Zhuofu Chen, William C. Feng, Aidan Burn, Junseok Park, Chong Chu, Hui Shen, Ting Wang, Alexander E. Urban, Xiaowei Zhu, Heng Li, Kathleen H. Burns, Hye-Jung E. Chun, Peter J. Park, SMaHT MEI Working Group, Alan P. Boyle, Ryan E. Mills, Weichen Zhou, Eunjung Alice Lee

**Affiliations:** Division of Genetics and Genomics, Boston Children’s Hospital, Boston, MA, USA; Department of Pediatrics, Harvard Medical School, Boston, MA, USA; Gilbert S. Omenn Department of Computational Medicine and Bioinformatics, University of Michigan Medical School, Ann Arbor, MI, USA; College of Informatics, Huazhong Agricultural University, Hongshan District, Wuhan, China; Department of Biomedical Informatics, Harvard Medical School, Boston, MA, USA; Department of Genetics, Washington University School of Medicine, St. Louis, MO, USA; Department of Neuroscience, City University of Hong Kong, Hong Kong SAR, China; Department of Pathology, Dana-Farber Cancer Institute and Harvard Medical School, Boston, MA, USA; Department of Epigenetics, Van Andel Research Institute, Grand Rapids, MI, USA; The Edison Family Center for Genome Sciences and Systems Biology, Washington University School of Medicine, St. Louis, MO, USA; McDonnell Genome Institute, Washington University School of Medicine, St. Louis, MO, USA; Department of Psychiatry and Behavioral Sciences, Stanford University, Stanford, CA, USA; Department of Genetics, Stanford University, Stanford, CA, USA; Department of Data Science, Dana-Farber Cancer Institute and Harvard Medical School, Boston, MA, USA; Broad Institute of MIT and Harvard, Boston, MA, USA; Ludwig Center at Harvard, Harvard Medical School, Boston, MA, USA; Department of Human Genetics, University of Michigan Medical School, Ann Arbor, MI, USA

**Author notes:** Correspondence (E.A.L.), (W.Z), (R.E.M), (A.P.B). These authors contributed equally (Seunghyun Wang, Mingyun Bae, Jinhao Wang).

## Abstract

Mobile element insertions (MEI) shape the human genome in both germline and somatic tissues. While inherited MEIs are well characterized, mapping somatic MEIs (sMEI) in non-cancer tissues remains challenging due to their low allelic fraction and repetitive nature. We established an integrative framework for sMEI analysis leveraging modern sequencing technologies and analytical innovations. We first benchmarked sMEI detection and demonstrated advantages of long-read and MEI-targeted sequencing for ultra-low-frequency events using a mixture of well-established cell lines. We then showed that haplotype phasing and donor-specific assemblies refine sMEI detection, effectively distinguishing from germline and false signals in *in-silico* tumor-normal mixtures. We further developed a source-tracing strategy based on internal sequence variation, expanding the catalogue of active source elements beyond traditional transduction-based methods. Applying this framework to donor tissues, we identified 18 rare somatic L1 insertions, revealing structural and source diversity. Our work provides a foundational framework and biological insight into sMEIs.

## Introduction

Mobile elements are a major source of structural variation in the human germline, yet their activity within the human body remains unknown. Mobile element insertions (MEIs) are generated through an RNA-mediated insertion mechanism known as target-primed reverse transcription (TPRT)^1,2^. In humans, only LINE-1 (L1) elements encode the proteins required for retrotransposition, enabling mobilization of not only L1s themselves but also other transposon-derived RNAs such as *Alu* and SVA, as well as processed mRNAs^3^. Because their insertions endanger genomic integrity^4–6^, induce genetic and epigenetic alterations^7–9^, and trigger immune responses^10^, mobile elements are kept under strict host surveillance through multiple layers of defense mechanisms, including epigenetic silencing of source elements and interfering RNAs^11^. Nevertheless, some elements evade this surveillance and contribute to disease initiation and progression, such as in cancers^6,12–14^ and neurological disorders^15–17^. Recent studies further indicate that somatic MEIs occur even in non-diseased normal human tissues^16,18–23^ and aging processes^11,24^, raising important questions about its prevalence and biological impact throughout the body.

Identification of sMEIs and their parental source elements in tissues remains challenging, with most obstacles rooted in current short-read sequencing technologies and low variant allele frequency (VAF). For instance, correct read alignment in repetitive genomic regions, such as segmental duplications^25,26^, is particularly problematic with short reads and hampers accurate variant calling. The repetitive nature of MEIs themselves further complicates the delineation of true somatic insertions when they arise within or adjacent to existing MEIs of nearly identical sequences (i.e., repeat-in-repeat)^27,28,29^. Short-read approaches are further limited by typically capturing only insertion junctions^30^, leaving complete insertion sequences unresolved, which often makes it difficult to confirm TPRT features and trance parental source elements. In cancer genomes, only 7.5-22.8% of sMEIs detected using short reads can be traced back to their sources, since source assignment relies on transduction sequences—unique flanking sequences occasionally co-mobilized with the source elements^13,31^. Because most sMEIs are present in only a small subset of cells within bulk tissue samples, their VAF is typically very low, which poses a significant challenge^32,33^. Several studies have reported sMEIs occurring in the human brain at such low VAFs, ranging from 0.09-0.4%^18^ or 0.53-1.2%^20^ estimated mosaicism. When reads are unevenly distributed between the two haplotypes, it becomes especially difficult to distinguish true heterozygous germline MEIs from somatic events.

Long-read whole genome sequencing (WGS) combined with analytical innovations provides opportunities to map sMEIs and gain biological insights into their epigenetic regulation as most platforms provide CpG methylation information^34,35^. Long reads that span large stretches of DNA—capturing entire insertions along with both junctions in a single molecule—can substantially improve sMEI detection. Yet even with long reads, reference bias persists where paralog content and sequence diverge from the sample^36^. Personalized, haplotype-resolved, donor-specific assemblies (DSA) overcome this limitation by accurately representing the individual’s alleles and copy number structure^37,38^. Aligning long reads directly to DSA provides a subject-matched coordinate system that normalizes germline divergence in variant calling, reduces misalignment across paralogous loci, and improves read placement through segmental duplications and other complex repeats. In particular, long reads link heterozygous germline variants and candidate sMEIs on the same molecule, enabling read-backed haplotype phasing against these assemblies^39^; this phasing information allows low-VAF signals to be correctly assigned to proper haplotypes and distinguished from noise. Recent studies have begun exploring the use of DSA to phase read alignment and improve somatic mutation discovery in limited samples, such as cancer and individual brain tissues^40,41^. However, the application of DSAs to somatic discovery, especially for sMEIs within complex regions, and systematic evaluations across different samples are still lacking. Long-read-based MEI analysis also enables direct profiling of internal repeat sequence variations, which may allow more comprehensive tracing of a parental source loci without relying on transductions^42^.

For cost-effective MEI profiling, several targeted approaches have been developed^16,22,43,44,45^. These methods enrich DNA templates containing MEIs and their genomic flanks, followed by either short-^16,22,43,44^ or long-read sequencing^45^, thereby enabling sensitive and scalable discovery of rare insertion events. Together, these advances establish numerous advantages of applying modern sequencing technologies and analytical approaches to investigate the ongoing mutagenesis in the human body by sMEIs. However, an integrative framework to fully leverage these technologies through systematic benchmarking is still lacking.

Here, we propose a multi-platform integrative framework for accurate sMEI detection and source tracing, and demonstrate its applicability in normal human tissues from the Somatic Mosaicism across Human Tissues (SMaHT) Network^46^. We first weigh the strength and the weakness of multiple sequencing platforms using a mixture of six highly characterized HapMap cell lines. For this, we report a high-confidence benchmarking callset for L1, *Alu*, and SVA, stratified into tiers based on the annotated TPRT features, and evaluate the detection performance of seven WGS-based computational methods and two MEI-targeted sequencing methods. We then propose an integration strategy whereby both insertion calls and raw signals from individual reads are combined for improved detection accuracy. We further demonstrate that haplotype phasing and DSA enable more accurate refinement of sMEI candidates and that our new source tracing method based on internal L1 sequence variations reveals haplotype-specific source L1 activities using *in silico* tumor-normal mixture experiments. We validate the applicability of our framework in normal tissue samples (liver, lung, colon, and brain) from four *post-mortem* human donors, and characterize sMEIs and their source diversity, including the first discovery of a somatic chimeric insertion involving small nuclear RNA and L1HS. Together, our work provides a practical framework for accurate sMEI mapping and expands our understanding of sMEIs in normal human tissues.

## Results

### Benchmark design for sMEI detection in HapMap mixture

Our aim is to establish a practical framework for sMEI analysis. As a first step, we sought to systematically benchmark sMEI detection approaches across sequencing platforms, depth, genomic context, and VAF levels. To enable such comparisons and to assess the strengths and the limitations of each platform and variant-calling method, we generated a high-confidence sMEI benchmarking set (**Table S1-S3**). Specifically, we identified non-reference MEIs with TPRT signatures from six HapMap cell lines (HG005, HG02622, HG02486, HG02257, HG002, and HG00438) and mimicked sMEIs in tissue mosaicism by mixing these cell lines using titres of varying frequency (**Figure 1A** and **S1A**) (The SMaHT network et al. in prep). In this HapMap mixture design, non-reference MEIs present in HG005 (83.5%, backbone genome) represent germline insertions, whereas those present in the other five cell lines but absent in HG005 represent sMEIs (**Figure S1B**). The benchmarking set was stratified into three tiers by annotating MEI subfamilies and TPRT features, such as poly-A tail and target site duplication (TSD), ranging from the most conservative (tier 1) to the most permissive (tier 3) (**Methods** and **Figure S1A**). The number of insertions in the benchmarking set varied by the class of mobile elements and their respective tiers, with *Alu* (n = 2,298; 2,431; and 2,552 in tiers 1–3, respectively) showing the highest counts, followed by L1 (n=408, 467, and 534) and SVA (n=175, 180, and 641) (**Table S4 and Figure S2**). The VAF, calculated by the mixing ratios (0.5% to 10%) and genotypes in individual cell lines, ranged from 0.25% to 16.5% across all three elements (‘expected VAF’; **Methods** and **Figure S2D**).

**Figure 1.**
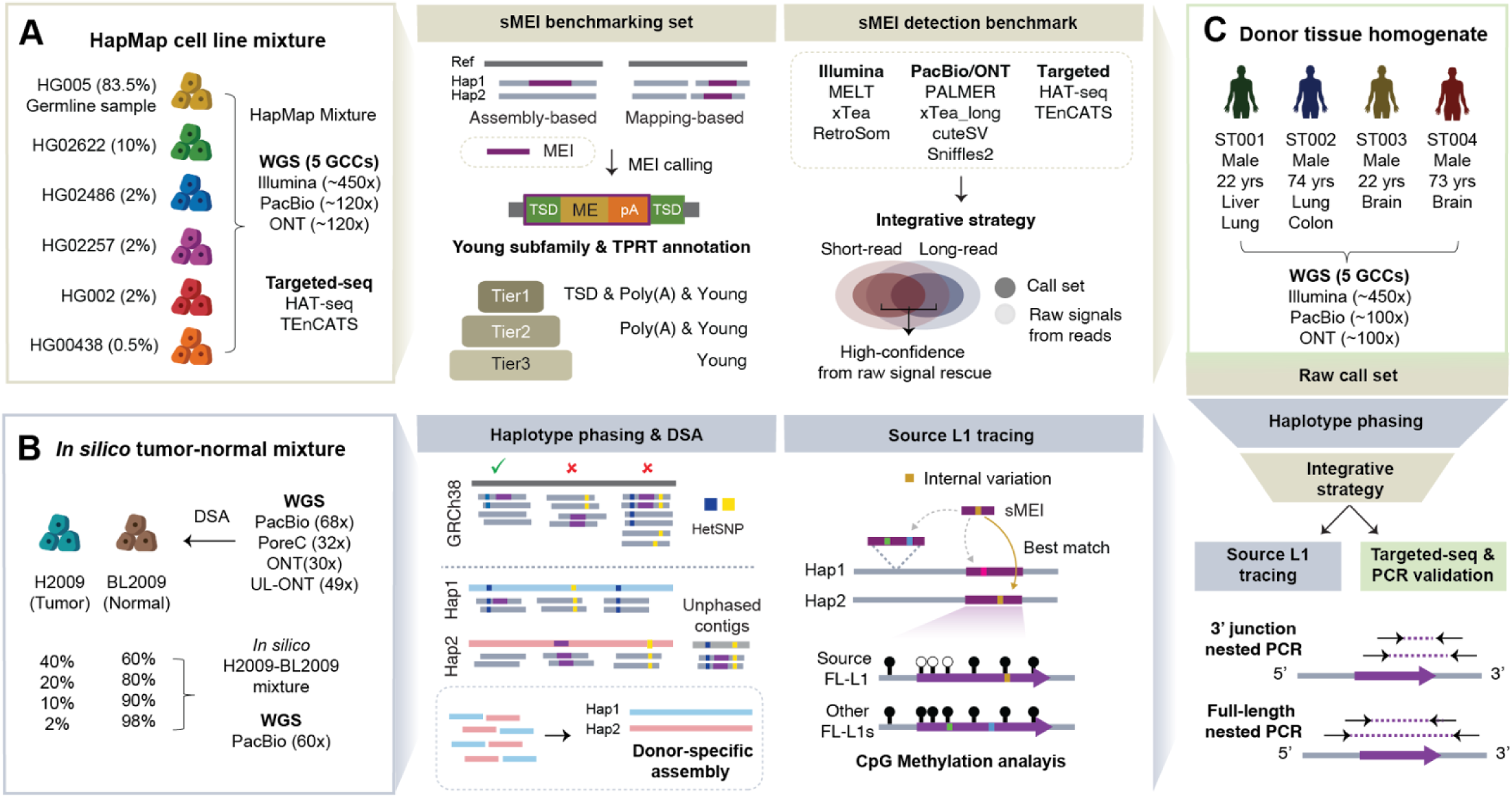
Overview of integrative framework for mapping sMEIs. **A**, Benchmarking sMEI detection methods using HapMap mixture from the SMaHT network. A benchmarking set was generated from PacBio and haplotype assemblies and stratified into three tiers by MEI subfamily and TPRT features. Nine sMEI detection methods were evaluated across sequencing platforms, depths, call set concordance, genomic regions, and VAFs. Integrative strategy incorporates both calls and raw signals from the reads to expand high-confidence sMEI candidates. GCC stands for Genome Characterization Centres in the SMaHT network. **B,** Refinement of sMEI detection using haplotype phasing and donor-specific assembly (DSA) and tracing of source L1s using internal sequence variation. In silico tumor-normal mixture (CASTLE project) dataset was generated to simulate sMEIs at various VAFs. Source L1s were traced using internal L1 sequence variation and their CpG methylation status was analyzed. FL-L1: full-length L1 **C,** The discovery of sMEIs in normal human tissues by applying the integrative framework for sMEI detection analysis. Each step established from HapMap mixture (sMEI detection method benchmark and multi-platform integration) and tumor-normal mixture (haplotype phasing/DSA and source L1 tracing) were integrated. The final sMEI candidates were confirmed by MEI targeted sequencing and nested PCR.

Most existing MEI detection tools were developed for identifying germline insertions or highly clonal sMEIs in cancer^30^ and their performance when applied to sMEIs in normal tissue has not been evaluated. We selected nine sMEI detection approaches, including seven WGS-based computational methods (**Methods**) and two MEI-targeted sequencing methods for benchmark. For methods originally developed for germline or highly clonal events (xTea_long^29^, cuteSV^47^ and Sniffles2^48^), we adjusted parameters by lowering supporting read counts; for general SV callers (cuteSV and Sniffles2), we additionally applied RepeatMasker^49^ to identify sMEI candidates among their insertion calls. For the WGS-based methods, we used high-depth bulk WGS data of the HapMap mixture provided by SMaHT network, including 450x short-read (Illumina), 110x long-read (PacBio and ONT) and their down sampling data (**Table S5**). Two MEI-targeted sequencing assays, HAT-seq^16^ and TEnCATS^45^ were applied to the genomic DNA of the HapMap mixture as orthogonal technologies (**Methods**). We report our results mainly focusing on L1 with the middle-tier (tier2) benchmarking sets, as L1s are the primary drivers of sMEI in somatic tissue, confirmed by our donor tissue analysis in this study. The results in the most conservative benchmarking set tier (tier1, **Figure S3**) and other element types (*Alu* and SVA, **Figure S4**) are provided in the Supplementary Information.

### Long-read and MEI-targeted sequencing improves sMEI detection at low VAFs and in challenging genomic contexts

We first evaluated the detection performance across sequencing platforms and depths (50x-432x Illumina, 30x-110x PacBio and ONT) using the benchmarking sMEI set (**Figure 2A and Table S6**). In the short-read data, we observed that the overall performance (F1 score) plateaued at ∼200x (xTea_mosaic and MELT), marking it as the recommended depth that balances cost and performance. The performance peaked at 300x, followed by a modest decline beyond this depth such that the difference in performance between 200x and 300x was subtle (<0.04 in F1 score). In the long-read data, such F1 scores were achievable at a much lower depth of ∼60x. For instance, xTea_mosaic showed the best performance (F1 score=0.53) at 200x Illumina among short-read WGS-based methods, a level matched by PALMER using only 60x PacBio (F1 score=0.59). The recall particularly improved from 30x to 60x across the long-read WGS-based methods (13.5% and 16.6% on average in PacBio and ONT, respectively). At the same depth, PacBio yielded from 15.3% to 24.1% higher F1 scores than ONT across the sequencing depth on average. Both short-read (HAT-seq) and long-read (TEnCATS) MEI-targeted sequencing methods showed higher performance than any WGS-based performance.

**Figure 2.**
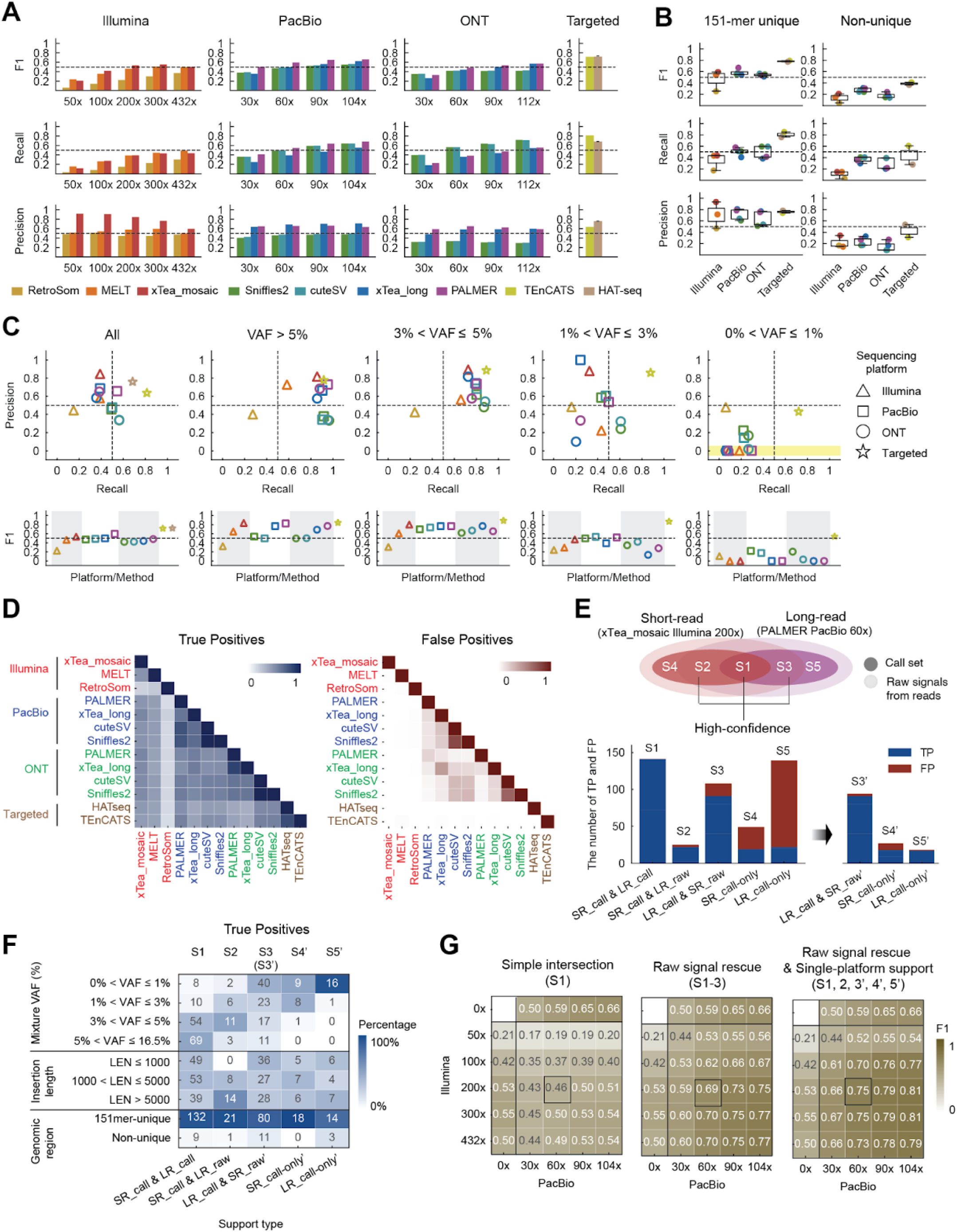
sMEI detection method benchmark and multi-platform (short and long read) integration using HapMap mixture. **A**, F1 scores, recall, and precision across sequencing platforms (Illumina, PacBio, ONT, and MEI-targeted sequencing) at various sequencing depths (full coverage WGS from BCM GCC and their downsampling data). HAT-seq performance is the average of four experimental replicates and the error bar was calculated based on the standard deviation. **B,** F1 scores, recall, and precision across 151-mer unique vs. non-unique regions. **C,** F1 scores, recall, and precision in each VAF bin. Platforms are indicated by shape. The dots located in the yellow area in the recall-precision scatter plot indicate that the corresponding method does not report the calls in the corresponding VAF bin. For HAT-seq, only recall is shown, because caller-level VAFs are not available for precision calculation. **A-C**, the same color key represents detection methods. **D,** Call set similarity (Jaccard index) of true positives (TPs) and false positives (FPs). Sequencing platforms are grouped by the same color. **B-D**, 200x and 60x callsets were used for short-read (Illumina) and long-read (PacBio and ONT) data, respectively. **E,** Schematic diagram of integrative strategy combined with raw signal rescues and the number of TP and FPs by each support type (S1-5) before and after FP filtering (S3’-5’). Support types are noted in the x-axis. **F,** Percentages and counts of true positives (TPs) across VAF bins, insertion length bins, and genomic regions, shown for each support type. **G,** F1 scores of multi-platform integrated callsets across different Illumina and PacBio coverage combinations.

We next examined whether the detection performance varied across genomic regions (**Figure 2B**) and used 200x Illumina and 60x PacBio/ONT as the representative sequencing depths (**Figure 2B-D**). We used a list of pre-defined, sample-agnostic genomic easy-to-map regions based on 151-mer uniqueness in pan-genome assemblies^50^. In our benchmarking set, 84.2% sMEIs fell into the 151-unique region (**Table S4**). The recall dropped sharply in the non-unique regions for all short-read WGS methods (10.8% on average), whereas long-read WGS-based methods maintained substantially higher recall (3.38-fold in PacBio and 2.79-fold in ONT) than short-read methods. These results strongly demonstrate that improved mappability with long reads broadens the callable genomic regions for sMEI.

Lastly, we assessed detection performance across different VAF ranges (**Figure 2C**). Detection patterns in variants with VAF >3% were largely consistent with those observed in the overall callset. At the VAFs lower than 3%, however, long-read WGS-based methods and MEI-targeted sequencing methods outperformed short-read methods, particularly in recall. When the VAFs reach extremely low (≤1%), long-read-based methods achieved 1.7-fold higher recall than short-read-based methods on average. This observation was consistent in a different benchmarking set (**Figure S3**) and for *Alu* and SVA (**Figure S4** and **Table S7-8**). This underscores the critical role of long-read sequencing in detecting low-frequency insertions in the normal human tissues. The MEI-targeted sequencing showed even higher recall in this VAF range (HAT-seq: 61.7%, TEnCATS: 72.0%), demonstrating cost-effective alternative to WGS-based approaches for detecting ultra-low VAF sMEIs.

### Integrating callsets with raw read-alignment signals enhances high-confidence sMEI identification

To explore an effective integration strategy across the callsets, we evaluated the concordance of the true positive (TP) and false positive (FP) calls between the methods (**Figure 2D**). Except RetroSom which show relatively low recall (**Figure 2A**), combinations of two callsets show 37.1-98.5% TP concordance. In contrast, FP concordance was markedly lower between the callsets derived from different platforms (0-2.0%), suggesting that cross-platform integration can effectively filter out FPs. Indeed, simple intersection of short- and long-read callsets (S1) achieved near-perfect precision with no overlapping FPs but reduced recall (recall=0.30) (**Figure 2E and S5**). Notably, 87.2% of calls in this intersection set had VAF above 3%, indicating that low-VAF calls and those with insufficient supporting reads were frequently missed (**Figure 2F**).

To mitigate the recall loss, we leveraged raw read-level signals to rescue calls detected by one platform but missed by the other, matching insertion sequences at the same breakpoint (**Methods**). This rescue strategy (S1-S3 in **Figure 2E**) improved performance 1.50-fold (F1=0.69, **Figure 2G**), with more TPs supported by both platforms (recall=0.54), while maintaining high precision of 0.98 (**Figure S5**). Notably, the rescued set captured insertions across distinct VAF levels, insertion lengths, and genomic regions (**Figure 2F**). For instance, only 9.1% of TPs called by short reads and rescued by long-read alignment signals (S2) showed VAF≤1%, whereas 44.0% of TPs called by long read and rescued by short-reads alignment signals (S3) showed VAF≤1%. Additionally, the largest number of sMEIs in non-unique regions were called by long reads with short-read alignment signals (S3). In contrast, more long insertions (>5Kbp) were called by short reads with long-read alignment signals (S2), reflecting the strength of short reads in detecting 3’ and 5’ insertion junctions regardless of insertion length. These patterns highlight that the multi-platform integration combined with raw read-level rescue enable to expand high-confidence sMEIs, complementing the limitation of each platform.

We further examined single-platform-only calls (S4 and S5) to reduce FPs by designing additional filtering schemes (**Figure S6**). For short-read-based xTea_mosaic calls, many FPs originated from the non-unique region (**Figure 2C and S6A**), and restricting the calls to the unique region removed 70% of FPs with nearly no recall loss (only one TP excluded) (S4’ in **Figure 2E**). For PALMER, two filters were effective: the old subfamily filter removed 82.1% of FPs without excluding any TP, while the sole-supplementary alignment filter removed 48.0% of young subfamily FPs. Together, these filtered out 99.1% of FPs with only 2 TPs lost (S5’ in **Figure 2E** and **S6C-D**). Notably, 93.8% of S5’ TPs were ultra-low mosaic sMEIs with VAF≤1% (**Figure 2G**). The same PALMER filters applied to the rescued callset S3 removed 82.4% of FPs (S3’). This enhanced integration callsets (S1-S2, S3’-S5’ in **Figure 2E**) achieved 1.63-fold improved detection performance (F1=0.75, recall=0.62 and precision=0.95) at Illumina 200x and PacBio 60x, compared to that of simple intersection between the two callsets (S1) (**Figure 2G**). When considering all combinations of Illumina and PacBio sequencing depths, the performances were 2.01-fold better than that of the simple intersections on average (**Figure 2G**). This improvement was also observed in the combination of Illumina and ONT (**Figure S7**). The performance of our integrative strategy was further validated using independent short-read and long-read WGS data (**Figure S8**).

### Phasing and DSA information increases precision in sMEI detection

Accurate read mapping is essential for alignment-based variant detection; however, repetitive genomic regions such as segmental duplications remain particularly challenging. For instance, inter-individual differences in segmental duplication copy number can cause reads originating from extra copies (absent in the reference genome) to misalign to existing loci, leading to false-positive somatic variant calls with low VAFs^25^. These artifacts can be effectively mitigated using long-read sequencing in combination with haplotype phasing and donor-specific assembly (DSA, **Figure 3A**), which provides improved resolution in complex genomic regions^41^. The application of phasing and DSA to HapMap mixtures is inherently limited due to their synthetic nature, where multiple genomes are mixed. To address this, we generated *in silico* tumor-normal (H2009-BL2009) mixtures from the CASTLE project^51^, enabling the evaluation of phasing and DSA as refinement strategies for sMEIs detection.

**Figure 3.**
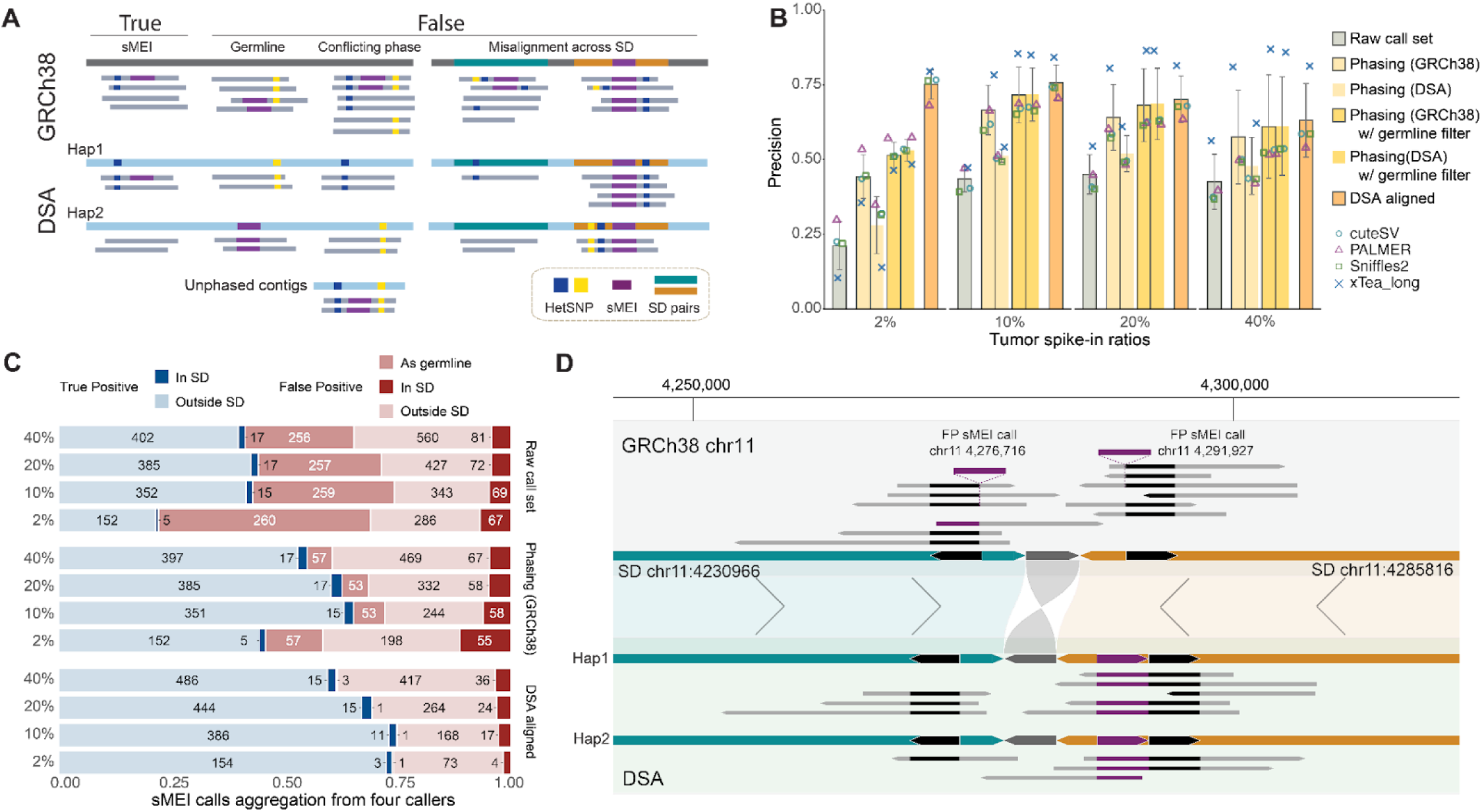
Refinement of sMEI detection from haplotype phasing and DSA analysis. **A**, Schematic illustrating how phasing information and DSA are leveraged to mitigate errors caused by germline MEIs, conflicting signals, or segmental duplications (SDs). **B**, Precision of sMEI calls across four tumor spike-in mixtures (2%, 10%, 20% and 40%). Bars indicate precision across four detection methods (cuteSV, PALMER, Sniffles2, xTea_long) using GRCh38-based initial alignment, DSA-based alignment, and various refinement methods (phasing on GRCh38, phasing on DSA, GRCh38-phasing with germline site filter, and DSA-phasing with germline site filter), coloured points show per-caller values. **C,** Composition of true positives (TPs) and false positives (FPs) inside vs. outside SDs, and FPs due to germline signals in the population, aggregated across four callers. Results are shown for GRCh38-based alignment, phasing on GRCh38, and DSA-based alignment, based on unique observations. **D,** Mis-mapping in a SD pair generates false sMEI calls on GRCh38 but not on the DSA. Upper panel: GRCh38 shows two 4.96kb reference blocks with L1 signal (black) within the inverted SD pair (chr11:4,230,966, green, and chr11:4,285,816, orange). Reads carrying an additional nearly identical 4.96kb L1 sequence (purple) align arbitrarily into the two blocks, yielding two apparent sMEIs at chr11:4,276,716 and chr11:4,291,927. Lower panel: In the DSA, a germline tandem duplicate (purple) was constructed in one SD; reads align consistently to both SD regions with no somatic signals.

We first generated a high-quality haplotype-resolved DSA using the BL2009 normal cell line data (**Methods** and **Figure S9**). Four mixtures of samples were prepared by spiking tumor sequencing data (PacBio) into normal at rates of 40%, 20%, 10%, and 2% to simulate varying frequencies of sMEIs. To evaluate the performance of sMEI discovery in these mixtures, we followed the similar pipeline as for the HapMap mixture above to generate a cancer benchmarking set of sMEIs in tumor H2009 cell line (L1 only, n=321, 441, and 448 in tier1-3, respectively) and excluding calls present in normal cell line data (**Methods** and **Table S9-S10**). The sMEIs in the cancer benchmarking set of these mixtures followed the expected distribution of VAF ranging from 1% to 20 % (**Figure S10A**). We next evaluated the performance improvement from the phasing and direct sequence assembly in three ways: (a) phasing information extracted from the sequencing reads aligned to the GRCh38 reference genome; (b) phasing information derived from the assembly contigs aligned to GRCh38; and (c) calling efficacy directly from the sequencing reads aligned to the DSA (**Figure 1B and 3A**). We further applied a population-level germline MEI filter to these sets by excluding polymorphic insertions previously detected by large scale sequencing projects (**Methods**).

Using phasing information from long-read sequencing, we performed local, read-backed phasing (see **Methods**) around each candidate insertion as the first refinement method. Relative to the pre-refined calls, precision increased by 13.7% to 55% in the 40% tumor spike-in sample (**Figure 3B** and **Table S11**), removing an average of 43.3% of false positives across callers. For assembly-derived phasing as mentioned as b, we utilized the empirical decision boundary (**Figure S10** and **Methods**) established in a previous study^41^ to distinguish between false positives and potential true positives, achieving 98.6% recall rate within the cancer benchmarking set and yielding 5.3% gain to an average of 47.9% precision in the 40% tumor spike-in sample.

Applying the germline MEI filter further increased precision in the 40% mixture for both read-backed phasing and assembly-derived phasing by 3.5% and 13.4%, reaching mean precisions of 61.0% and 61.2%, respectively, while maintaining recalls up to 86.0% and 81.2%. We observed a similar pattern at other mixture levels. Notably, improvements extended into the most challenging, lowest mixture level sample: in the 2% spike-in, precision rose from 21.1% to 53.1%, highlighting the value of phasing information for ultra low-frequency sMEI detection (**Figure 3B** and **S11**).

We then asked whether aligning reads directly to an individually-matched DSA would further suppress reference-mismatch artifacts. Because a DSA intrinsically encodes the donor’s germline alleles and architecture, direct DSA alignment normalizes germline divergence and reduces mis-mapping in segmental duplications (SDs) and other complex genomic regions, thereby facilitating identification of *bona fide* somatic insertion signals absent from the DSA. Indeed, sMEI calls derived from DSA alignments demonstrated the highest precision among three refinement methods, ranging from an average of 63.1% in the 40% mixture to a remarkable 75.1% in the 2% mixture. We further systematically assessed true and false positive sMEIs in the context of SDs and known germline sites across the three refinement methods. Read-derived phasing methods shows a substantial refinement of sMEI signals (**Figure 3C**); For example in the 40% mixture, the aggregated number of unique FP calls across callers in SD decreased from 67 in the pre-refined sets to 55, while FPs in germline sites decreased from 256 to 57. Notably, calls from DSA-based alignment yielded sparse FPs in the SD regions, ranging from 36 (40% mixture) to 4 (2% mixture). Additionally, only 0.2% (15 calls in total) of DSA-aligned calls overlapped with predefined germline sites (**Methods**). We observed that DSA substantially reduced the proportion of SD-localized false positives (adjusted OR = 0.61, 95% CI 0.52-0.72; *p*=2.48x10^-9^; **Methods**), leading to a clear shift in call compositions across refinement methods (**Figure 3C**) while preserving true positives.

As a case study, we examined two potential false positive somatic L1 insertions that fell within SD regions as aligned to GRCh38: a 4.96kb insertion (A) at chr11:4,276,716 and another (B) of the same length at chr11:4,291,927, both concordantly detected by cuteSV and Sniffles2 in the 40% spike-in sample, with insertion (A) additionally detected by xTea_long (**Figure 3D and Figure S12**). One copy of the SD pair at chr11:4,230,966 and its inverted homologous copy at chr11:4,285,816 both contain a 4.96kb reference segment with L1 signal. This segment was confirmed to be 99% identical to the insertion sequences, supported by 11 of 46 reads in call A and 9 of 74 reads in call B, yielding two putative somatic L1 calls. However, in the haplotype-resolved DSA, we identified a germline homozygous signal in one SD pair (chr11:4,285,816), which represents a tandem duplication of the reference segment. When realigning the same molecules to the DSA, all supporting reads for both A and B were redistributed to the reconstructed SD (chr11:4,285,816 in GRCh38), which carries this germline tandem duplicate with L1 signal. In contrast, the other DSA SD copy (chr11:4,230,966 in GRCh38) showed no evidence of L1 insertion. This provides strong evidence that the DSA successfully reconstructed the exact insertion segment in both haplotypes, while in GRCh38 the reads were mis-mapped and arbitrarily assigned between these duplicated SD blocks (**Figure 3D**). This showcases how two false positive L1 insertions can be introduced by a homozygous MEI germline event on the same SD copy and illustrates how homologous SD pairs in the traditional human reference introduce ambiguous alignments, leading the supporting reads for a germline insertion in one SD to be misdistributed and misinterpreted as somatic MEIs at one or both loci. By reconstructing the germline insertion segment, the DSA facilitates accurate mapping and eliminates these false somatic signals.

### Internal L1 sequence variation enables haplotype-resolved source tracing

The human genome contains ∼100 full-length L1HS (FL-L1HS) loci that are retrotransposition-competent^52^. However, which FL-L1HS loci contribute to somatic retrotransposition in human tissues is still an open question. Traditionally, tracing somatic L1 insertions back to their source loci requires 3’ transduction sequences, leaving a substantial fraction of insertions untraceable^42^. Only 8.8%^53^ to 15%^27^ of germine L1 insertions contain transduction sequences, while for somatic L1 insertions, this number can range from 11% in normal colon tissue^21^ to 35% in gastrointestinal tumors^42^.

Long-read sequencing enables the profiling of complete L1 insertion sequences, giving an opportunity to use the internal L1 sequence for source tracing. To assess the feasibility of this approach, we first collected reference and non-reference FL-L1HS sequences from the BL2009 cell line, referred to as the FL-L1HS catalogue, totaling 627 unique sequences (566 reference and 61 non-reference; **Methods and Table S12**). Next, using a transduction-based tracing approach, we established an evaluation set of 59 insertions confidently linked to 17 source loci. These source sequences showed considerable sequence diversity, suggesting a possibility of source tracing using the sequence variation (**Figure S13A**). We developed a source-tracing pipeline that leverages L1 internal sequence variation compared against the FL-L1HS catalogue (**Figure 4A** and **Methods**). Applied to the transduction-based evaluation set, our pipeline successfully mapped 25 of 59 insertions (42.4% recall) to their sources with 100% precision (**Figure 4A**).

**Figure 4.**
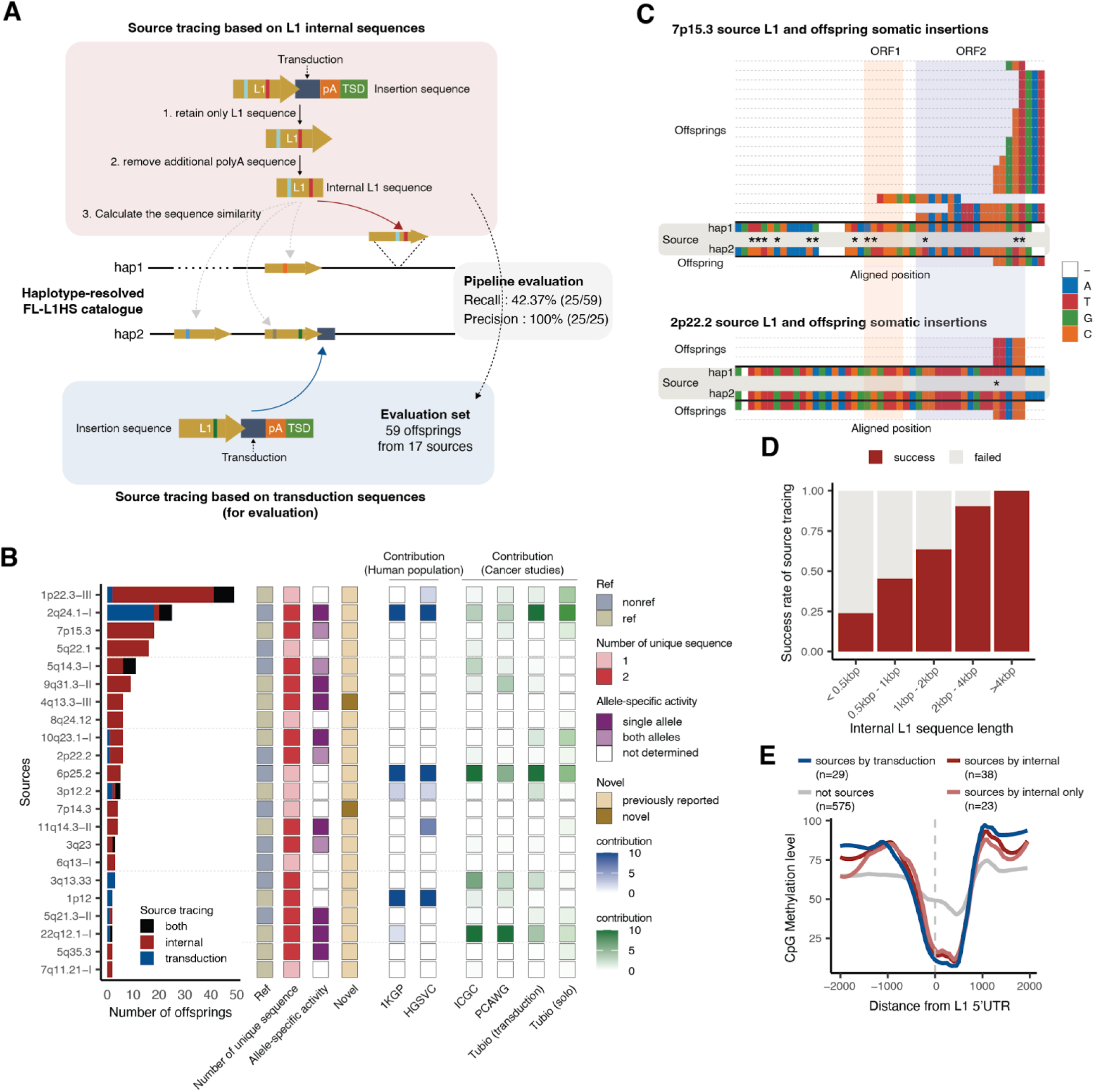
L1 source tracing using internal sequence variations in tumor-normal mixture. **A**, Schematic overview of the transduction-based and internal sequence-based source tracing pipelines. Recall and precision was shown for the internal sequence-based source tracing method. **B**, Comprehensive characterization of L1 source loci with at least two offspring. The left bar chart shows the number of offspring from each source, with color indicating the prediction method (transduction, internal sequence, or both). The adjacent columns provide detailed annotations for each source locus, including reference (ref) or non-reference (nonref) status, unique allelic sequences, allele-specific activity, and source activity reported in human population studies (1KGP^54^, HGSVC^28^) and cancer (ICGC^31^, PCAWG^55^, Tubio^42^) studies. For allele-specific activity, sources detected only by the transduction method and containing only one unique allelic sequence are shown in white, as allele-specific activity cannot be calculated for them. For other sources inferred from L1 internal sequence variation, if two unique allelic sequences are present at the source locus, allele-specific activity was calculated and was colored according to the major haplotype. **C**, Examples of two sources (7p15.3 and 2p22.2) and their offspring. Multiple sequence alignment of offspring sequences and source sequences was performed using Clustal Omega^56^. Bases differing among the four source sequences were extracted, with asterisks indicating haplotype-specific variants. **D**, Success rate of source tracing using L1HS internal sequence variations depending on the internal L1 sequence size. **E**, DNA methylation level at the 5’UTR (±2kb) of FL-L1HS for four source categories of source loci in the H2009 tumor cell line.

We then applied the pipeline to 448 somatic L1 insertions identified from the H2009 cancer WGS and successfully traced 172 (38.3%) insertions to 38 unique source loci (**Table S13**). Compared to the transduction-based method alone, our approach traced 2.9-fold more insertions to 2.2-fold more source L1s (**Figure S13B**). However, 34 (57.6%) of 59 insertions with 3’ transduction could only be traced by their transduction sequences, indicating that the two strategies are complementary and should be combined for comprehensive source tracing. Using both approaches, we traced sources for 206 (46%) insertions in total (**Figure 4B**). As expected, sources identified by transduction were often previously reported, whereas several identified by L1 internal sequence variation were novel (**Figure 4B**).

Our pipeline can reveal haplotype-specific source activities in homozygous FL-L1HS loci. We calculated the allele-specific contributions of source loci with two unique sequences using the internal sequence variations. Notably, source L1 transcripts are often derived from one haplotype, suggesting that both allele-specific sequence variation and heterogeneous epigenetic status may influence the allelic activity of source L1 element. (**Figure 4B**). For example, a source L1 element at the 7p15.3 locus produced the highest number of somatic insertions, but with a strong allelic bias (**Figure 4C**). By contrast, a source L1 at the 2p22.2 locus contributed insertions equally from both alleles (**Figure 4C**). The two haplotypes differed by 12 base pairs at 7p15.3 and by only one base pair at 2p22.2. Since our pipeline relies on internal sequence variation, its performance improves with increasing offspring insertion length. For somatic insertions longer than 2 kbp, we were able to successfully trace 90% of events back to their source loci (**Figure 4D and S13C**).

We further examined haplotype-specific CpG methylation at the promoters (5’ UTR) of the identified source L1s. In the H2009 tumor cell line, sources identified by our pipeline exhibited significant hypomethylation. Notably, sources identified exclusively by internal sequence variation showed similar hypomethylation to transduction-based sources, reflecting their active epigenetic state (**Figure 4E**). We found the promoters of these same sources were hypermethylated in the normal cell line (BL2009), confirming their cancer-specific active state (**Figure S13D**). Overall, our source tracing pipeline leveraging L1 sequence diversity significantly expands the number of identifiable source loci.

### A multi-platform integrative framework reveals structural and source diversity of low-frequency sMEIs in normal human tissues

Finally, we assessed the applicability of our integrative framework (haplotype phasing, multi-platform integration, and source L1 tracing) for sMEI analysis by analyzing six tissue samples—liver, lung, colon, and brain—from four *post-mortem* human donors (ST001-ST004; **Figure 1C**) (The SMaHT network et al. in prep). Each step and resulting callset in our framework was illustrated using WGS data of the ST003 brain sample, which have similar sequencing depths to what we used as the representative dataset in the HapMap mixture (**Figure 5A** and **Table S14**). We identified a total of 18 somatic L1 insertions with variable frequency across samples (**Figure 5B** and **Table S15**). Each sample had at least one high-confidence insertion supported by both platforms—called by long reads and rescued by short reads (S3’, **Figure S15**). Twelve additional L1 insertion candidates were supported exclusively by long reads only (S5’) and passed manual inspection. Given their estimated low VAFs (only one supporting reads detected; **Table S15**), this result coincided with the HapMap mixture benchmark, which showed that low VAF insertions were primarily supported by long reads (**Figure 2F**).

**Figure 5.**
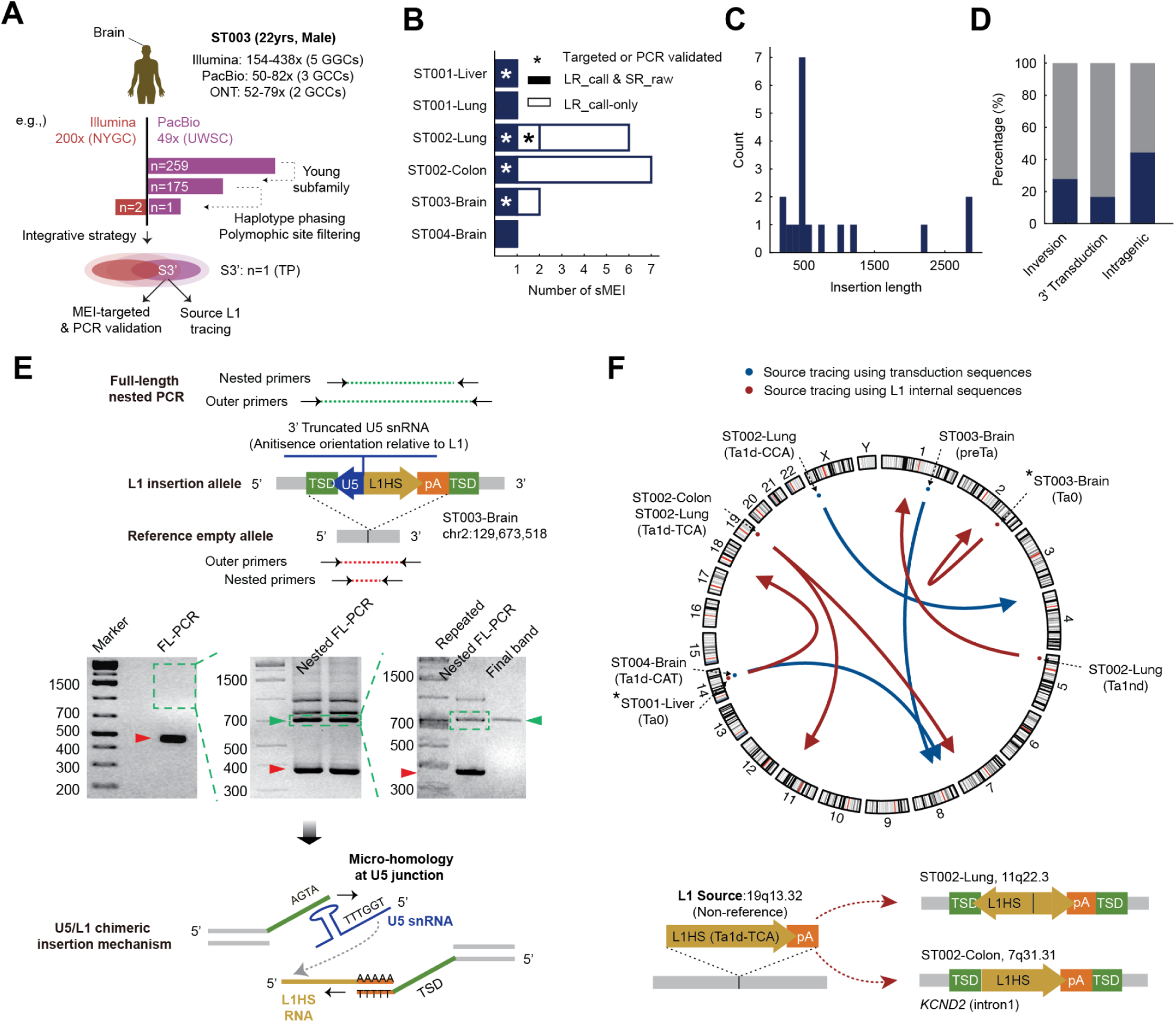
Characterization of sMEIs and source L1s in normal human tissue homogenates. **A**, Overview example of the somatic L1 detection pipeline applied to ST003-brain yielding one high-confidence insertion supported by long-read call and short-read alignment signal. This process was repeated for all samples from each of 5 GCCs and for other donors. **B,** The barplot shows the number of somatic L1 insertions per sample. **C,** Length distribution of somatic L1 insertions across the tissue samples. **D** Somatic L1 categorization by 5’ inversion, 3’ transduction, and intragenic vs. intergenic genomic regions. **E,** Schematic diagram of the U5/L1 chimeric insertion and representative agarose gels (bottom) for the insertion in ST003 brain sample. Due to low mosaicism, the L1 insertion band was not visible after the initial PCR (FL-PCR). A region of the gel at the expected size was excised (dashed green box) for subsequent DNA extraction and re-amplification. This repeated nested PCR approach successfully isolated a clean band corresponding to the L1 insertion allele (green arrowheads). Red arrowheads indicate the band from the reference empty allele. **F,** Source L1s identified by transduction and internal L1 sequences and schematic diagram of the L1 insertions in ST002 lung and colon samples from the same L1 source traced by internal L1 sequence variation.

Re-sampling and validating somatic insertions from bulk tissue are challenging due to their low VAFs and likely focal distribution. Nonetheless, we performed the nested PCR for eight selected candidates by amplifying the 3’ L1-genome junctions (**Table S16**), using either the same gDNA extracted from the tissue homogenate that was used to generate the corresponding WGS data (ST001 and ST002) or the gDNA extracted from a different core but in the same tissue (all four donors). Three candidates with estimated VAF larger than 0.5% (ST001 liver, ST002 colon and ST003 brain) were validated in both cases (**Figure 5B, S15** and **Table S15**). By contrast, the two insertions in ST002 lung with lower estimated VAF (0.20-0.25%) were only validated in the same gDNA extracted from the tissue homogenate and two other insertions in ST002 lung with similar estimated VAF (0.25%) or much lower (below 0.20%) were not validated even in the same gDNA extracted from the tissue homogenate (**Table S15**). Among the five insertions validated by 3’ nested PCR, three insertions in ST001 liver, ST002 colon, and ST003 brain samples were also detected by TEnCATS (**Figure S16**), while two out of these three insertions (ST002 colon and ST003 brain samples) were also detected by HAT-seq, all performed on different cores from the homogenate used for WGS.

All identified insertions were 3′-truncated (72.2%, 13/18 shorter than 1 kbp), with no full-length insertions observed (**Figure 5C**). Intra-inversion, indicative of twin-priming, was observed in 27.8% (5/18) of the insertions (**Figure 5D**). It is lower than that reported in pan-cancer analyses (40%)^13^ consistent with observations from normal colorectal epithelium (29.5%)^21^, but relatively higher than the rate (21% to 25%) reported in the cell culture or population studies^27,53,57^. This suggests that twin-priming rates may vary across tissue types and biological contexts. Interestingly, we found one somatic L1 insertion in a brain sample whose sequence contained a 3’-truncated U5 snRNA (*RNU5A-1*) fused in an antisense orientation to a 5’-truncated L1HS element, a structural pattern consistent with very rare polymorphic events previously reported^58–61^ (**Figure 5E**). The micro-homology found at the U5-L1 and insertion site junctions is consistent with a twin-priming model of formation^60^. We validated its sequence and structure using nested full-length PCR followed by Sanger sequencing (**Figure 5E** and **S15**). To our knowledge, this is the first report of a somatic snRNA/L1 chimeric insertion in human tissues, and long-read-based analysis will enable more comprehensive investigation of these non-canonical forms of sMEI in the somatic state.

Our source tracing pipeline identified eight somatic insertions originating from seven source L1s (**Figure 5F**). Three insertions (16.7%) with 3′ transductions originated from two reference and one non-reference full-length source L1s. The other five insertions were traced to their source loci based on internal sequence variation. The identified source L1s belonged to various L1HS subfamilies: preTa, Ta0, Ta1nd, Ta1d-CCA, Tad1d-CAT, and Ta1d-TCA (**Figure 5F**). Notably, two somatic L1insertions found in the lung and the colon samples of the same donor (ST002) were derived from the same source of Ta1d-TCA subfamily, known active L1HS subfamily^62^ (**Figure 5F**). Eight (44.4%) somatic insertions were located within introns of seven different protein coding genes, including genes associated with tissue-specific diseases. For example, one insertion in ST001 liver occurred in intron 14 of *PTPRM* , a gene involved in cell adhesion^63^, liver fibrosis^64^ , hepatocellular carcinoma^65^. Another insertion in ST004 bain occurred in intron 2 of *CSMD1* , implicated in neurodevelopmental or psychiatric disorders such as schizophrenia^66,67^ and intellectual disability^68^. Most somatic insertions detected in normal tissues are likely neutral, but further investigation of their clonal structure across tissues and their functional impact may provide insights into their regulation and potential roles in altering the epigenetic and transcriptomic landscape in the human body.

## Discussions

Somatic mobile element insertions are increasingly recognized as contributors to genomic diversity in normal tissues, yet their reliable detection has remained challenging due to their low frequency, repetitive nature, and the lack of standardized benchmarks. To overcome these challenges, we established an integrative framework for sMEI analysis and demonstrated its applicability in normal human tissues. As a first step of our framework, we weighed the strengths and weaknesses of multiple sequencing platforms using a mixture of the six highly characterized HapMap cell lines. We generated the sMEI benchmarking set stratified by TPRT features and evaluated nine detection methods across platforms, depth, genomic stratification and VAFs. The detection performance at 200x sequencing depth using short reads was achievable at much lower 60x using long reads.

Our benchmarking analysis highlighted the value of long-read and MEI-targeted sequencing for detecting sMEIs in difficult genomic regions and at low VAFs. The higher detection performance of MEI-targeted sequencing at ultra-low VAFs suggests that it is a cost-effective complement to WGS-based sMEI detection. Furthermore, we established a cross-platform integration strategy that leverages read-level raw signals with additional filtering schemes accounting for platform-specific errors. This approach enhanced the high-confidence sMEI detections supported by both platforms, maintaining high precision. Overall, our systematic evaluation of sMEI detection across sequencing depths and platforms provides practical guidance for effective study design.

Using a tumor-normal mixture sample, we further confirmed that the haplotype phasing and DSA alignment enable the refinements of sMEI signals by filtering germline and false somatic signals. DSA contiguity is a key limitation of our approach and varies substantially across tissues and donors. Using ONT and 10x Genomics linked-reads data, bridging and scaffolding raised phased N50 in a neurotypical dorsolateral prefrontal cortex from 0.75 Mb to 2.63 Mb without Hi-C or trio information^41^, yet in the benchmarking donor ST001 (liver and lung) contiguity increased only from 0.45 Mb to 0.72 Mb. Such variability, likely driven by DNA integrity, extraction, library preparation differences, and tissue-specific fragmentation, constrain haplotype block lengths. While these assemblies are sufficient for sMEI discovery and refinement, shorter contigs increase the risk of phase switches and mis-joins, reduce the callable fraction in complex regions, and may limit confident resolution of copy number variations and other megabase-scale variants. Consequently, interpretations should be explicitly conditioned on each sample’s phased N50 and effective haplotype span.

Leveraging long-read technology, we demonstrated that full-length L1 sequences harbor sufficient sequence variation to serve as reliable source markers and developed a new source-tracing pipeline based on the internal L1 sequence variation. Our pipeline identified three times more sources compared to the traditional transduction-based method. It is particularly effective in normal tissues, where transductions are expected to be less common . Although our method substantially expands source identification, it has several limitations. The accuracy of this approach depends heavily on the quality and completeness of the full length L1HS reference catalogue. While most source elements belong to the L1HS subfamily, rare trans-subfamily insertions may be missed by this pipeline. In addition, its performance with ONT data has not yet been fully evaluated. Future work will aim to address these limitations and further refine the pipeline into a more comprehensive and robust framework for L1 source tracing.

Our integrative framework enabled reliable detection of low-frequency mosaic L1 insertions in normal human tissues, revealing structural and source diversity. Future studies that integrate single-cell and spatial sequencing as well as functional assays will be essential to determine how sMEIs influence cellular function. Analyzing larger and more diverse donor cohorts will clarify the prevalence, tissue distribution, and inter-individual variability of sMEIs and their active source loci. Additional questions also remain: whether the sMEI landscape in normal tissues mirrors the high variability observed across cancer types, whether specific genetic factors influence sMEI frequencies, and whether distinct sets of source loci or hotspot loci are active across different tissues during development and aging. Our integrative framework lays the foundation for answering these questions and advancing our understanding of the role of MEIs in human health.

## Methods

### Dataset for WGS-based methods

#### HapMap mixture

Briefly, the SMaHT network designed HapMap mixtures by combining six cell lines from International HapMap Projects at predefined ratios to simulate somatic variants: HG005 (83.5%), HG02622 (10%), HG02486 (2%), HG02257 (2%), HG002 (2%), and HG00438 (0.5%). Each of the five SMaHT GCCs generated high-coverage WGS data for the mixtures, including short-read (Illumina) and long-read (PacBio and ONT) datasets. We obtained the aligned sequencing data (BAM) of the mixtures from the SMaHT data portal (https://data.smaht.org/) for benchmarking WGS-based computational sMEI detection methods.

#### In silico BL2009-H2009 mixtures

PacBio-HIFI sequencing data for BL2009 (normal) and H2009 (lung adenocarcinoma) cell lines were obtained from the Sequence Read Archive (SRA, PRJNA1086849) and the CASTLE project (https://github.com/CASTLE-Panel/castle)^69^. We generated four *in silico* BL2009-H2009 mixtures at 60x coverage with tumor read spike-in ratios of 2%, 10%, 20%, and 40%, respectively. Ideally, the heterozygous sMEIs in the H2009 tumor cell line will be downsampled to the predefined VAFs (1%, 5%, 10%, and 20%). The mixed reads were aligned to the GRCh38 human reference genome using pbmm2 (https://github.com/PacificBiosciences/pbmm2).

#### Human tissue homogenate

The SMaHT network collected six tissue samples from four *postmortem* donors: ST001 (liver and lung), ST002 (lung and colon), ST003 (brain), and ST004 (brain). To ensure cellular homogeneity, the tissue samples were homogenized prior to sequencing. WGS was performed by five SMaHT GCCs using Illumina, PacBio, and ONT platforms. Aligned sequencing data (BAM) of each sample were obtained from the SMaHT data portal (https://data.smaht.org/).

Detailed sample preparation and processing protocols for the sample that we obtained from the SMaHT network are described in the SMaHT flagship publication (The SMaHT network et al. in prep). Sequencing depth information for each sample is provided in **Table S4**.

### Library preparation and sequencing in MEI targeted sequencing

#### TEnCATS

TEnCATS library preparation was performed following McDonald et. al.^45^ with changes described here for SQK-LSK114 and R10.4.1 flow cells (**Code availability**). 30µL of gDNA from the HapMap cell mixture was dephosphorylated in a 40μL reaction with 6μL Quick CIP (M0525S, NEB) and 4μL 10X rCutSmart buffer (B7204S, NEB). This reaction was inverted and gently tapped to mix, and then incubated at 37°C for 30 minutes followed by a 2 minute heat inactivation at 80°C. The Cas9 ribonucleoprotein (RNP) was formed by combining 850ng of in vitro transcribed guide RNA, 1µL of a 1:5 dilution of Alt-R S.p. HiFi Cas9 Nuclease V3 (1081060, IDT), and 1X rCutSmart buffer (B7204S, NEB) in a total of 30µL. This reaction was incubated at RT for 20 minutes. Next, both the prepped gDNA and RNP were placed on ice and the RNP was added to the dephosphorylated gDNA. 1μL 10mM dATP and 1.5μL Taq DNA Polymerase (M0273S, NEB) were added to the gDNA:RNP reaction, then inverted and gently tapped to mix. This reaction was incubated at 37°C for 30 minutes for Cas9 cutting and brought to 75°C for a-tailing for 10 minutes. For adapter ligation, the cut reaction was transferred to a 1.5mL microcentrifuge tube. We then added 5μL T4 DNA ligase (M0202M, NEB) and 5μL Ligation Adapter (LA; SQK-LSK114, ONT). This reaction was inverted to mix and incubated at RT for 20 minutes with rotation. Following ligation, we added 1 volume of 1X Tris EDTA (TE) and inverted to mix. Next 0.4X Ampure beads (SQK-LSK114, ONT) are added and incubated for 5 minutes with rotation followed by 5 minutes at RT without rotation. The beads were then washed twice with 150μL Long Fragment Buffer (LFB; SQK-LSK114, ONT) followed by incubation with 50μL Elution Buffer (EB; SQK-LSK114, ONT) at 37°C for 30 minutes. Finally, we loaded the R10.4.1 MinION flow cell following the ONT protocol using 12μL of the library and sequenced for 72 hrs on a MinION. For sequencing of the libraries on PromethION flow cells, 12μL of the same library sequenced on a MinION was used and brought to a total volume of 32μL with EB then sequenced for 72 hours on a PromethION2 Solo.

#### HAT-seq

We prepared Human Active Transposon-Sequencing (HAT-seq) libraries by adapting the protocol from previous work^16^ . Starting with 500 ng of genomic DNA extracted from the SMaHT HapMap cell line mixture, we first fragmented the DNA using a Covaris R230 focused-ultrasonicator (Covaris, LLC). These fragments were subsequently processed for end repair, A-tailing, and adapter ligation with the KAPA HyperPrep Kit (Kapa Biosystems, Inc). Using the adapter-ligated DNA as a template, we generated four technical replicate libraries. The L1 enrichment and indexing PCR steps included two key modifications to the previous protocol: 1) the P7_Ns_L1Hs primer contained a mix of 2,4,6,8 staggers to enhance the sequence diversity of the semi-amplicon library. 2) unique dual indices were used for both P5 and P7 adapters to ensure precise library demultiplexing. Library fragments ranging from 340 to 450 bp were enriched using a dual size selection with AMPure beads from 0.575x to 0.75x beads ratio (Beckman Coulter). We quantified libraries with the KAPA Library Quantification Kit (KAPA Biosystems) and then combined them into an equimolar pool. The final library pool was sequenced with a 5% PhiX spike-in on an Illumina NovaSeq sequencing system, generating 2x150 bp paired-end reads. A list of adapter and primer sequences used for HAT-seq library construction is available in **Table S17**.

### Donor-specific assembly generation

For cancer cell line mixture analysis, we built donor-specific assemblies (DSA) using PacBio-HIFI, Nanopore ONT, UL-ONT, and Pore-C sequencing data from the normal cell line (BL2009) in the CASTLE panel project (SRA PRJNA1086849)^69^. These data were collected to generate a DSA without downsampling for this normal-tissue pair using Verkko^70^ v2.2.1, the resulting DSA then went through a decontamination process through FCS v0.5.5 (**Figure S9**. Tumor DSA construct).

For benchmarking donors, due to the lack of phasing information such as Hi-C or trio data, we defined the *target* as the draft donor-specific assembly and introduced the concept of *bridges*—additional sequencing datasets that share overlap with the target to extend phase blocks within the DSA (Supplementary Figure. Donor DSA construct). The initial draft DSA was assembled from PacBio HiFi, Oxford Nanopore (ONT), and HERRO-corrected ONT reads. HiFi reads were randomly downsampled to 100× coverage, and ONT reads were corrected with HERRO^71^ using the *model_R10_v0.1.pt* model, retaining only reads longer than 10 kb. The draft DSA was partitioned into two haplotypes using a custom graph-based approach: contig overlaps were used to construct a weighted graph, and a greedy bipartitioning algorithm assigned contigs to haplotypes so as to maximize phase block continuity within each group. Phased heterozygous SNVs were then identified using dipcall^72^ v0.3 on the two resulting haplotypes. For bridges, original ONT reads were used (rather than HERRO-corrected reads), as HERRO correction discards base quality scores. Phased heterozygous SNVs were called using PEPPER-Margin-DeepVariant r0.8.

Draft DSA contigs (“targets”) were connected via ONT reads (“bridges”) using overlapping phased heterozygous SNPs as anchors. For each merged interval, phased genotypes at each shared SNP between target and bridge blocks were compared, and the most common phase switch decision (match or flip) was determined for each target phase block. A concordance matrix was constructed, and target blocks connected through bridges with valid phased SNPs overlap were grouped using a disjoint-set union (DSU) algorithm. Within each group, genomic coordinates were merged to define extended phase blocks. The flip status for each phase block was propagated from an anchor block, ensuring consistent haplotype assignment across the group. Whenever a flip was detected, contigs previously assigned to haplotype 1 were switched to haplotype 2 and vice versa, maintaining phase consistency until another flip was encountered. Final phase switch corrected phase blocks were exported in BED format for downstream analysis. Phase switch-corrected contigs were further scaffolded using RagTag v2.1.0 and gap-filled with ONT reads using TGS-Gapcloser v1.2.1. For each haplotype, only reads assigned to the corresponding haplotype were used in the scaffolding and gap-filling steps. The quality assessment of all diploid assemblies was performed thoroughly by three tools: BUSCO^73^ v5.7.1, QUAST^74^ v5.2.0, and Merqury^75^ v.1.3 (Supplement Table. DSA stats).

### Benchmarking set generation

#### HapMap mixtures

To simulate somatic insertions in HapMap mixtures, we first identified high-confidence non-reference MEIs in six HapMap cell lines (**Figure S1A**). We obtained publicly available PacBio long-read whole-genome sequencing data (BAM files) and haplotype-resolved assemblies for these cell lines from the Human Pangenome Reference Consortium (humanpangenome.org)^39^. We called germline insertions using a combination of assembly-based methods (DipCall v0.20^40^, Minigraph v0.34^41^) and an alignment-based method (Sniffles v2.2^30^).

After calling the insertions using three different callers, we extracted insertions annotated to any repeat type using RepeatMasker, which database was merged with both Dfam^76^ and RepBase v.20181026. This process yielded a set of candidate mobile element insertions. For more detailed repeat annotation, we used BLAST against consensus sequences from RepeatBrowser^77^, To focus on potentially active elements, we restricted our analysis to insertions from young subfamilies known to be active in the human genome: L1HS and L1PA2 for L1 family; *Alu*Y, *Alu*Yb8, and *Alu*Ya5 for *Alu* family; and SVA_E and SVA_F for SVA family.

We then developed a custom pipeline to identify the canonical signatures of target-primed reverse transcription: target site duplications (TSD) and polyA/T tails. To identify TSD sequences, we generated synthetic paired-end reads for each insertion. For each event, we extracted 500bp of the upstream and downstream reference sequence to serve as the left and right pseudo-read, respectively. Then, the first 100bp of the insertion sequences was appended to the left pseudo-read, and the last 100bp of the insertion sequence was prepended to the right pseudo-reads. These synthetic reads were then aligned to the reference genome using bwa-mem with default options, and the resulting CIGAR strings were inspected. A TSD was identified if the alignment match (“M” in the CIGAR string) extended beyond the original 500bp reference sequence length. This method allowed us to annotate the TSD sequence and its size, defining a canonical TSD as being between 4 and 30bp in length. polyA/T tails were identified by searching for sequences of more than 10 A at the 3’ end or 10T at the 5’ end of the insertion, allowing for up to two mismatches after excluding the TSD sequence.

Based on the subfamily, TSD, polyA/T features, we stratified the filtered insertions into three confidence tiers. Tier 1 included insertions from young subfamilies with canonical TPRT signatures: both a TSD and a polyA/T. Tier2 included insertions with only a polyA tail including events with target site deletions, and Tier 3 comprised all remaining young subfamily insertions, regardless of TPRT evidence. To annotate zygosity, we calculated the VAF for each insertion and applied a Gaussian Mixture Model to the resulting distribution to distinguish between heterozygous and homozygous events. Finally, we manually inspected all candidate insertions by comparing them with another manually curated set of mobile element insertions in the HG002 cell line from a previous study^29^, and then applied the same pipeline to the other five cell lines.

After establishing a high-confidence set of MEIs for each of the six HapMap cell lines, we combined all identified insertions to simulate the somatic insertions in the HapMap mixture. For this simulation, any insertion present in the HG005 cell line was assigned as germline, while insertions found only in the other cell lines were assigned as somatic. The expected VAF in the cell line mixture was then calculated using the following equation:

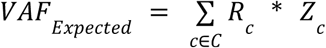

where *C* is the set of cell lines {HG005, HG02622, HG02486, HG02257, HG00, HG00438}, *R_c_* is the mixture ratio for cell line *c*, and *Z_c_* is the zygosity of the insertion in cell line *c* (0.5 for heterozygous, 1.0 for homozygous) (**Figure S1B**). For tiering of HapMap mixture insertion, we assigned the most frequently occurring tier for the final benchmarking set (**Table S1-S3**).

#### In silico BL2009/H2009 mixtures

The H2009 lung cancer cell line exhibits extensive copy number variation, which prevents the generation of a diploid assembly from long-read sequencing data. Therefore, instead of using assembly-based methods, we applied multiple read-based structural variation callers (nanomonsv v0.7.0^76,78^, Severus v1.2^51^, and Sniffles2 v2.2^48^) in a paired tumor-normal mode with the matched normal BL2009 cell line. The resulting insertion calls were processed using the same annotation pipeline as the HapMap cell lines, including subfamily classification, TSD identification, and polyA/T tail analysis. As the majority of detected insertions belonged to the L1HS subfamily, we constructed a benchmarking set focused exclusively on L1 elements. All insertions, particularly those reported by only a single caller, were manually inspected, and confidence tiers were assigned using the same criteria as the HapMap mixture benchmarking set (**Table S9**).

### sMEI detection

The WGS-based methods were included if they are capable of detecting sMEIs in single-sample mode given normal human tissues where matched case–control samples are not available and meet one of the following criteria: (1) specifically designed for sMEI detection (xTea_mosaic^29^ and RetroSom^20^), (2) used for sMEI detection in previous studies^41,79^ (MELT^54^ and PALMER^27^), or (3) originally developed for germline MEIs but adaptable for sMEI detection with simple parameter tuning—lowering supporting read counts (xTea_long^29^, cuteSV^47^, and Snilffles2^48^) (**Code availability**). Since cuteSV and Sniffles2 are general structural variant callers, we applied RepeatMasker^49^ to annotate sMEI candidates for their insertion calls.

#### xTea_mosaic

We ran a mosaic mode (-M) of xTea^29^ (v0.1.9) with the modified parameters related to the number of supporting clipped and discordant reads for sMEI detection at the lower frequency. We lowered the cutoffs for the minimum number of clipped reads (--nclip 2, default 3), clipped reads whose mates map in the repetitive region (--cr 0, default 1), clipped reads in the filtering step (--nflip 1, default 3), and discordant pair in the filtering step (--nfdisc 1, default 5).

#### MELT

Mate coordinates and insert size field tags were added to BAM files using samtools^80^ (fixmate) to improve the runtime of MELT. MELT^54^(v2.2.2) was implemented with default parameters in ‘Single’ genome analysis mode.

#### RetroSom

RetroSom^20^ (v2) recommends to run control and case samples independently (step 1-6) and combine the results for somatic insertion detection (step 7-8). However, since our study aims to detect sMEI in the normal tissues without matched control-case samples, we ran RetroSoms as single sample mode by running step 1-6 using only HapMap mixture samples. The maximum number of supporting reads to be considered as a putative somatic insertion (-n) was set as 30% of sequencing depth of the sample. We omitted the polymorphic filtering step since the sMEI in the HapMap mixture originated from the germline insertions.

#### PALMER

PALMER^27,28,41,45^ (v2.0.1) was implemented with the default parameter setting. The calls with at least one and two high-confidence supporting reads were included in the callsets of PacBio-HIFI and ONT sequences, respectively. PALMER then implements a masking process to exclude the calls in low-confidence regions for the final callsets. The mask file can be found here https://doi.org/10.5281/zenodo.1573339081.

#### xTea_long

Since xTea_long^29^ (v0.1.0) was originally developed for germline insertion detection, we modified the parameters related to the number of supporting reads to detect sMEIs (--user -c 1) in the clip (-C) and assembly (-A) step.

#### cuteSV

cuteSV^47^ (v2.1.2) was implemented with a modified parameter for the number of supporting reads for sMEI detection (--min_support 1, default 10). Since cuteSV was originally developed for SV detection, we applied RepeatMasker and remained the calls pass following three criteria: (1) ‘L1’, ‘*Alu*’, ‘SVA’ class, (2) less than <10% divergence (combined % of substitution, deletion, and insertion bp) by comparing consensus ME sequence, (3) insertion sequence larger than 50bp.

#### Sniffles2

We ran two different modes of Sniffles2 (v.2.6.3.)^48^. We first ran a germline mode of Sniffles2 with a modified parameter for the number of supporting reads for sMEI detection (--minsupport 1). Additionally, we ran a mosaic mode of Sniffles2 for sMEI detection (--mosaic). Since the minimum VAF levels of mosaic mode was set as 5%, we lowered the threshold of minimum VAF levels to also detect the sMEI lower than 5% (--mosaic-af-min 0, default 0.05). Then, we applied RepeatMasker and the same filtering criteria with cuteSV. The performance from modifying the number of supporting reads was mainly used for the evaluation and that of mosaic mode was provided in **Table S5-7.**

#### TEnCATS

For both L1 and *Alu* datasets, on-target rate, and MEI calling were characterized using NanoPal (**Code availability**), adapted from our prior study^45^. Briefly, reads were aligned to the reference genome and non-reference TE insertions were detected with PALMER. Next, TEnCATS reads were then classified into on-target using BLASTn and reads supporting the MEIs are clustered by location. Variant calls with fewer than two read support were filtered from the final results.

#### HAT-seq

Bulk HAT-seq data were analyzed using a computational pipeline for the SMaHT HapMap cell line mixture, adapted from our prior study^16^. Raw, de-multiplexed FASTQ reads were trimmed of adapters and low-quality bases. Read pairs containing L1Hs sequences in Read 2 were retained; the corresponding Read 1, capturing the 3′ genomic flank, was aligned to the human reference genome (build hg38/GRCh38, GCA_000001405.15, without alternate contigs). Putative L1 insertion sites were then identified by peak calling.

Identified peaks were annotated by comparison to insertion databases and with their genomic context and quantitative features (*i.e.* height in reads per million [RPM], width, template count). Peaks were subsequently classified as known reference (KR), known non-reference (KNR), unknown (UNK)/somatic insertions, or artifacts. For the artificial KNR L1 spike-ins designed to mimic somatic events in HapMap mixture, a rescue step was implemented: after initial classification as KNRs, they were rescued and re-routed to undergo the same downstream filtering to generate the final unknown (UNK)/somatic insertion callset.

### Performance evaluation

To evaluate the performance of each sMEI detection method, we computed recall, precision and F1 score of each call set against the benchmarking set. Using the germline insertion information (present in the HG005) in the benchmarking set, germline calls were excluded from the callsets to ensure that evaluation was limited to somatic candidates. A call was classified as a true positive (TP) if its breakpoint was located within ±100 bp (for WGS-based methods) or ±200 bp (MEI-targeted sequencing) of an insertion in the benchmarking set; otherwise, it was considered a false positive (FP).

Recall was defined as the proportion of true sMEIs in the benchmarking set that were successfully detected by the method:

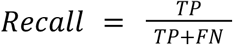

Precision was defined as the proportion of detected insertions that were true sMEIs. To avoid penalizing calls that are absent in the current benchmarking set tier but present in a lower-confidence tier, we masked such calls during precision calculation. For example, when evaluating a call set against the tier 2 benchmarking set, we excluded calls that would be considered true positives in the tier 3 set from the false positive count. Precision was then calculated as:

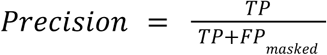

F1 score, the harmonic mean of recall and precision, was calculated as:

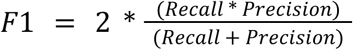

All metrics were evaluated globally as well as in stratified analyses by genomic regions and VAF. Specifically, to evaluate the performance in each genomic region, we used two genomic stratifications: (1) 151-mer unique vs. non-unique region and (2) ME-free vs. ME region. We obtained genomic stratification bed files from the SMaHT data portal (https://data.smaht.org/) and used the combination of ‘easy’ and ‘difficult’ region as 151-mer unique and ‘extreme’ region as non-unique region, respectively. We defined the ME region based on ‘L1’, ‘Alu’, and ‘SVA’ annotation in the reference genome by RepeatMasker^49^. The benchmarking sets and the callsets then were stratified by genomic regions based on the breakpoint positions. Recall and precision were then calculated independently within each region using the same equation as in the global analysis:

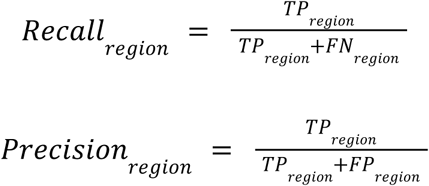

Similarly, to evaluate the performance in each VAF range, we used different VAF definitions in recall and precision calculation. In recall, the benchmarking set was stratified into four VAF bins (0%<VAF≤1%, 1%<VAF≤3%, 3%<VAF≤5%, and VAF>5%), based on the ‘expected VAF’ that we defined in the benchmarking set. Each VAF bin was designed to include enough insertions to ensure robustness of the performance (**Table S4**). The entire call set was then compared to the stratified benchmarking set in each VAF bin to calculate recall:

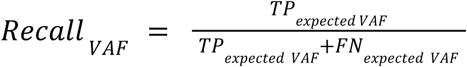

On the other hand, for precision, we used ‘caller VAF’ that each method reports for each call, since the mixture VAF for false positives cannot be defined in the benchmarking set. So, the call set was stratified into same VAF bins with the recall using the caller VAF and the entire benchmarking set was compared to the stratified call set to calculate precision:

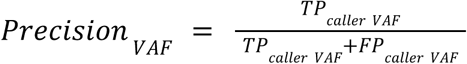

#### Integrative strategy combined with call set and raw read alignment signals

We selected xTea_mosaic and PALMER for the integration—two methods that showed the best overall performance in benchmarking in each platform. The supporting type of each call in the integrated call set was classified into five types according to short-read call (SR_call), short-read raw signal (SR_raw), long-read call set (LR_call), and long-read raw signal (LR_raw). In the integrated call set, we first included calls supported by both methods (SR_call & LR_call, S1) with overlapping insertion breakpoints (within 200bp). Except for the calls belonging to S1, we searched the raw signals in orthogonal platforms. To this end, we extracted the 7bp from the breakpoint coordinates in the clipped reads and examined whether this sequence matched with the insertion sequence identified in the other platform. The calls that have raw signals from the other platforms were classified as (SR_call & LR_raw (S2) or LR_call & SR_raw (S3)). For the calls supported by a single platform (SR_call-only (S4) and LR_call-only (S5)), we applied method-specific FP filtering strategies. For the xTea_mosaic set, we excluded the calls located in the non 151-mer unique regions (S4’). For the PALMER call set, we ran BLAST^82^ using the complete insertion sequence and only retained calls classified into young subfamilies (L1HS and L1PA2). We also excluded calls that were supported by only supplementary alignment reads (S5’). We also applied the same FP strategy to S3 (S3’). The final integrated callsets included the calls belonging to five supporting types: S1, S2, S3’, S4’ and S5’.

### Refinement of sMEIs using phasing information

To determine the sMEI, we performed local phasing using long-read sequencing data. First, we called single nucleotide polymorphisms (SNPs) in a 40 kbp window centered on the insertion locus using GATK4 HaplotypeCaller. From the resulting variants, we selected up to five high-quality (QUAL > 1000) heterozygous SNPs (hetSNPs) that were closest to the insertion site for the analysis. For phasing, we analyzed long reads that covered both a selected hetSNP locus and the corresponding insertion site. This phasing process was conducted for each of the selected hetSNPs (up to five per insertion), and the final classification was assigned based on the most frequent outcome. An insertion was categorized as ‘No hetSNPs’ if no suitable heterozygous SNPs were identified within the 40 kbp flanking region, or as ‘No covering reads’ if no single long read spanned both the insertion site and a selected hetSNP locus. If the insertion was linked to alleles from both haplotypes, it was classified as ‘Insertion in multiple haplotypes’. The insertion was classified as ‘Germline’ if it was confidently linked to a single haplotype while the alternate haplotype was well-covered by reads lacking the insertion. Finally, an insertion was labeled as ‘Somatic MEI (sMEI)’ if it was linked to a single haplotype but supported by a low number of reads, indicative of a potential somatic or mosaic event.

### Refinement of sMEIs using phasing information from DSA

We employed two strategies to leverage phasing information for refining somatic MEIs using DSA: an GRCh38 reference-based approach and a DSA reference-based approach. For the GRCh38-based approach, tumor-normal mixture data were aligned to the GRCh38 reference genome. Reads were then phased using LRphasing (**Code Availability**) based on phased heterozygous SNVs identified from the DSA, and phased VAFs were then computed for each haplotype. The haplotype with the higher phased VAF was designated as hapA, provided the two haplotypes were supported by at least five reads. To further reduce false-positive somatic sMEI candidates, we applied an empirical decision boundary defined by VAF_HapB_ = 0.2 × VAF_HapA_. Variants lying outside this boundary were flagged as probable artifacts and excluded from downstream analyses.

### DSA-based alignment analysis

Six samples were aligned to the DSA using pbmm2. To prevent alignment to unphased contigs, only contigs assigned to haplotype 1 and haplotype 2 were extracted and merged to create the reference genome. Callers (PALMER, cuteSV, Sniffles2 and xTea_long) were then run on each DSA-aligned sample with the DSA as the customized reference genome. For xTea_long, a DSA-specific repeat library was generated with the library preparation module of xTea (**Code Availability**).

We aligned DSA contigs to GRCh38 and performed coordinate projection between DSA and GRCh38 with a custom pipeline (**Code Availability**). A projection was declared successful only if the relevant contig-to-reference alignment had MAPQ ≥ 60 and fully covered the ±100 bp window around the queried coordinate. Sites lacking such coverage were excluded from rate denominators. The same criterion was applied symmetrically for DSA-to-GRCh38 projections. We did the projections for MEI calls and the population-polymorphic germline sites.

To evaluate whether DSA improves false positives (FPs) in segmental duplications (SD) more than outside SD. Let g = (caller, mixture, method), for each g, among FP calls, we counted

- 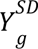 = # FP in SD
- 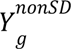 = # FP outside SD
- *n_g_* = 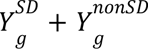

Polymorphic germline calls were excluded from the analysis.

We fit a binomial GLM with log link (logistic regression) to FP composition:

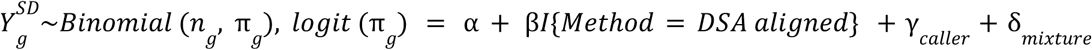

Caller and mixture level were included as fixed effects with 4 levels each. Coefficient β is the log odds ratio comparing DSA aligned calls to raw call set for “FP in SD vs. outside SD”, adjusted for caller and mixture. The adjusted odds ratio (OR) is *e*^β^.

We tested *H*_0_: β ≥ 0 vs *H*_1_: β < 0 using the Wald z statistic from GLM. We reported the adjusted OR with 95% confidence interval and the one-sided p-value. We also checked for overdispersion, where it indicated no overdispersion (residual deviance/df = 0.998, p = 0.984), supporting the binomial variance assumption. All analyses above were implemented in R (v4.4.1)^83^.

### Constructing the germline full-Length L1HS sequence catalogue in haplotype resolution

Germline L1HS elements can exist in both reference and non-reference forms. To extract all full-length reference L1HS sequences, we first identified candidate loci from RepeatMasker tracks and filtered out entries shorter than 4000 bp. Next, we extracted reads mapped to L1HS regions with mapping quality over 30, excluding supplementary alignments. These reads were converted into FASTA format and assembled using hifiasm^84^ with default parameters except for the `-f0` option. Unitig GFA files generated from the assembly were converted to FASTA, and RepeatMasker was used to annotate L1HS loci. The L1HS sequences were then extracted from these locally assembled FASTA files. If the L1HS sequences were identical between the two haplotypes, only one unitig was present in the FASTA file. Otherwise, two unitigs represented the distinct haplotypes.

Non-reference L1HS sequences were extracted using a similar approach, except for the identification of L1HS loci. When a non-reference L1HS insertion occurred at a locus overlapping a reference L1HS, RepeatMasker annotations alone were insufficient to identify the correct position. Therefore, we re-aligned the locally assembled FASTA files to the reference genome using pbmm2, and extracted the insertion sequences from the aligned assemblies.

After obtaining fully resolved L1HS sequences, we identified full-length elements via BLAST^82^ alignment against the 5′UTR, ORF1, and ORF2 sequences. The reference sequences for 5′UTR, ORF1, and ORF2 are provided in the supplementary materials.

### Source identification based on transduction sequences

To extract the transduction sequence from insertion sequence, we first performed a BLAST^82^ search using repeat consensus sequences. If either the left or right flanking sequence of an insertion was longer than 50 bp, we further aligned these flanking sequences to the human genome. If the flanking sequence mapped within 5000 bp of any full-length L1HS catalogue, whether reference or non-reference, we defined that locus as the source.

### Source inference based on internal L1HS sequences

During mobile element insertion, small structural variations such as insertions, deletions, or inversions can occur. Additional poly-A signals may also be appended during transcription. To avoid confounding elements, we extracted only the internal L1HS sequence based on BLAST^82^ annotation of the insertion sequence and further removed the polyA/T tails. The trimmed sequence was then aligned on the haplotype-resolved FL-L1HS catalogue using BLAST^82^, and the locus with the highest bitscore was identified as the putative source. If the bitscores of the top and second hits were identical, we annotated the insertion as “source untraceable”.

### Discovery of sMEIs in human tissue homogenates

We first ran xTea_mosaic (Illumina) and PALMER (PacBio and ONT) to generate callsets for each tissue WGS sample generated from each of five GCCs (**Table S4**). For the PALMER call set, MEI subfamily of each call was annotated using BLAST, and the old subfamily calls and supplementary alignment calls were proactively filtered out. Then, we applied haplotype phasing to filter out germline calls. To exclude the calls that arose from misalignment or misclassification during haplotype phasing, we cross-checked the haplotype phasing results across the SMaHT tissue samples and the calls classified as germline in at least one sample were also classified as germline in other samples. To further exclude germline calls, we curated the polymorphic MEI insertion positions from previous studies that used short-read^29,31,54,85–88^ and long-read^27,28,38^ sequencing and excluded overlapping sites (±100 bp).

Next, we applied the integrative strategy (‘Integrative strategy combined with call set and raw read-level signals’ in **Methods**) to all combinations of short-read and long-read callsets. For example, ST003 sample has five short-read and five (three PacBio and two ONT) long-read WGS samples (**Figure 5A**), and we performed the integrative strategies for 25 combinations. All candidates are manually inspected by at least two reviewers, by checking the genomic content (Gene, SegmentalDuplicate, and RapeatMasker tracks) and insertion information in the reads using IGV^89^ and BLAT^90^.

The VAF of each insertion was estimated by average the VAF in each sample which was calculated by dividing the number of supporting reads by the local depth in the long-read WGS data. The L1HS subfamily of source L1s was identified by canonical positions defining L1HS subfamilies from previous work^62^.

### Experimental PCR validation

Putative somatic L1 insertions were validated using two nested PCR strategies adapted from the previous work^16^. For 3′ junction validation, the first-round PCR (20 µL) contained 20 ng of gDNA, 10 µL of 2× KAPA2G Robust HotStart ReadyMix (KAPA Biosystems, KK5702), and 0.5 µL each of L1Hs-AC-28 and a site-specific 3′ primer (20 µM). Cycling conditions were: 95°C for 3 min; 20 cycles of 95°C for 15s, 60°C for 15s, and 72°C for 15s; and a final extension at 72°C for 1 min. For the second round, 1∼2 µL of the first-round product was re-amplified with L1Hs-nest-28 and a nested 3′ primer for 20, 25, or 30 cycles to optimize target band visibility.

For full-length PCR validation, the first-round PCR contained 50 ng of gDNA with 1 µL of PrimeSTAR GXL DNA Polymerase (Takara Bio, R050A), 5 μL of PrimeSTAR GXL buffer (5×), 2 µL of dNTPs (2.5 mM each), and 1 µL each of site-specific 5′ and 3′ primers. Cycling conditions were: 98°C for 3 min; 30 cycles of 98°C for 20 s, 60°C for 20 s, and 68°C for 1 min 15 s; and a final extension at 68°C for 3 min. The resulting full-length amplicon was excised from the gel, purified, and used as a template for a second amplification round with nested primers.

All amplicons were visualized on 2% agarose gels. The 3′ junction PCR products were submitted for Amplicon-EZ sequencing (Azenta, Inc.). Full-length L1 amplicons were excised and purified using the PureLink™ Quick Gel Extraction Kit (Invitrogen, K210012) and PCR Purification Kit (Invitrogen, K310001). A portion of the purified product was submitted for Sanger sequencing, while the remainder was sonicated into fragments (Covaris R230) for deep sequencing via Amplicon-EZ. Primer sequences used for validation are listed in **Table S16**.

## Data availability

This study was conducted as part of the NIH Common Fund consortium initiative, Somatic Mosaicism across Human Tissues (SMaHT). The benchmark datasets described in this study are available through dbGaP (http://www.ncbi.nlm.nih.gov/gap) under the study accession number, phs004193. The data used in this work was provided by the SMaHT Data Analysis Center (DAC) on behalf of the SMaHT network. More information about the SMaHT network is available online at https://smaht.org/, about the SMaHT Data Portal at https://data.smaht.org/, and types of data generated by the Network at https://data.smaht.org/about/consortium/data. The sequencing samples of H2009 and BL2009 can be found at CASTLE project (https://github.com/CASTLE-Panel/castle). The open access data (callsets for HapMap mixture and tumor-cancer mixture, and haplotype-resolved assembly for BL2009) is available at https://doi.org/10.5281/zenodo.1725434491. We used GRCh38 (GenBank accession: GCA_000001405.15) as the primary reference in this project. The reference can be downloaded at https://ftp.ncbi.nlm.nih.gov/genomes/all/GCA/000/001/405/GCA_000001405.15_GRCh38/seqs_for_alignment_ <u>pipelines.ucsc_ids/</u>.

## Code availability

The scripts, command lines, and related pipeline links used in this project can be found at https://github.com/ealee-lab/SMaHT-MEI-Benchmarking.

## Acknowledgments

We are extremely grateful to the SMaHT donors, and donor families, who have generously provided such precious gifts to support this important work. This research is supported by the NIH Common Fund, through the Office of Strategic Coordination/Office of the NIH Director under awards 1UM1DA058230, UM1DA058219, UG3NS132127, UG3NS132084 and UG3NS132146.

## Author information

E.A.L., W.Z., R.E.M., and A.P.B. supervised the project. M.B., X.Z., and Z.W. generated the benchmarking set. S.W., C.E.S., B.M., S.D., Z.C., J.P., C.C., T.W., A.E.U., X.Z., and W.Z. generated callsets for WGS-based computational methods. B.Z., K.N., S.M., J.S., S.J.L., T.L.M., C.M., and W.Z. generated callsets for MEI-targeted sequencing. S.W performed benchmarking and integrative strategy. S.W., M.B., J.W., H.L., and W.Z. designed and performed haplotype phasing and DSA analysis. M.B developed an internal sequence variation-based source tracing pipeline. B.Z,, K.N., S.M., R.K.G., and A.M.T. performed nested PCR validation. A.B., H.S., and K.H.B., prepared and managed samples for MEI-targeted sequencing and PCR validation. W.C.F. and H.E.C. managed sequencing data. S.W., M.B., J.W., B.Z., R.E.M., W.Z., and E.A.L drafted the manuscript. S.W., M.B., J.W., A.P.B., R.E.M., W.Z., and E.A.L revised the manuscript with input from all authors. All authors read and approved the final version.

## Conflict of interest

All other authors declare no conflict.

## Notes

### Competing Interest Statement

The authors have declared no competing interest.

## Reference

1. Feng, Q., Moran, J.V., Kazazian, H.H., Jr, and Boeke, J.D. (1996). Human L1 retrotransposon encodes a conserved endonuclease required for retrotransposition. Cell 87, 905–916.

2. Cost, G.J., Feng, Q., Jacquier, A., and Boeke, J.D. (2002). Human L1 element target-primed reverse transcription in vitro. EMBO J 21, 5899–5910.

3. Cordaux, R., and Batzer, M.A. (2009). The impact of retrotransposons on human genome evolution. Nat Rev Genet 10, 691–703.

4. Symer, D.E., Connelly, C., Szak, S.T., Caputo, E.M., Cost, G.J., Parmigiani, G., and Boeke, J.D. (2002). Human l1 retrotransposition is associated with genetic instability in vivo. Cell 110, 327–338.

5. Gilbert, N., Lutz-Prigge, S., and Moran, J.V. (2002). Genomic deletions created upon LINE-1 retrotransposition. Cell 110, 315–325.

6. Rodriguez-Martin, B., Alvarez, E.G., Baez-Ortega, A., Zamora, J., Supek, F., Demeulemeester, J., Santamarina, M., Ju, Y.S., Temes, J., Garcia-Souto, D., et al. (2020). Pan-cancer analysis of whole genomes identifies driver rearrangements promoted by LINE-1 retrotransposition. Nat Genet 52, 306–319.

7. Miki, Y., Nishisho, I., Horii, A., Miyoshi, Y., Utsunomiya, J., Kinzler, K.W., Vogelstein, B., and Nakamura, Y. (1992). Disruption of the APC gene by a retrotransposal insertion of L1 sequence in a colon cancer. Cancer Res 52, 643–645.

8. Scott, E.C., Gardner, E.J., Masood, A., Chuang, N.T., Vertino, P.M., and Devine, S.E. (2016). A hot L1 retrotransposon evades somatic repression and initiates human colorectal cancer. Genome Res 26, 745–755.

9. Slotkin, R.K., and Martienssen, R. (2007). Transposable elements and the epigenetic regulation of the genome. Nat Rev Genet 8, 272–285.

10. De Cecco, M., Ito, T., Petrashen, A.P., Elias, A.E., Skvir, N.J., Criscione, S.W., Caligiana, A., Brocculi, G., Adney, E.M., Boeke, J.D., et al. (2019). L1 drives IFN in senescent cells and promotes age-associated inflammation. Nature 566, 73–78.

11. Gorbunova, V., Seluanov, A., Mita, P., McKerrow, W., Fenyö, D., Boeke, J.D., Linker, S.B., Gage, F.H., Kreiling, J.A., Petrashen, A.P., et al. (2021). The role of retrotransposable elements in ageing and age-associated diseases. Nature 596, 43–53.

12. Lee, E., Iskow, R., Yang, L., Gokcumen, O., Haseley, P., Luquette, L.J., 3rd, Lohr, J.G., Harris, C.C., Ding, L., Wilson, R.K., et al. (2012). Landscape of somatic retrotransposition in human cancers. Science 337, 967–971.

13. Solovyov, A., Behr, J.M., Hoyos, D., Banks, E., Drong, A.W., Thornlow, B., Zhong, J.Z., Garcia-Rivera, E., McKerrow, W., Chu, C., et al. (2025). Pan-cancer multi-omic model of LINE-1 activity reveals locus heterogeneity of retrotransposition efficiency. Nat Commun 16, 2049.

14. Shukla, R., Upton, K.R., Muñoz-Lopez, M., Gerhardt, D.J., Fisher, M.E., Nguyen, T., Brennan, P.M., Baillie, J.K., Collino, A., Ghisletti, S., et al. (2013). Endogenous retrotransposition activates oncogenic pathways in hepatocellular carcinoma. Cell 153, 101–111.

15. Bundo, M., Toyoshima, M., Okada, Y., Akamatsu, W., Ueda, J., Nemoto-Miyauchi, T., Sunaga, F., Toritsuka, M., Ikawa, D., Kakita, A., et al. (2014). Increased l1 retrotransposition in the neuronal genome in schizophrenia. Neuron 81, 306–313.

16. Zhao, B., Wu, Q., Ye, A.Y., Guo, J., Zheng, X., Yang, X., Yan, L., Liu, Q.-R., Hyde, T.M., Wei, L., et al. (2019). Somatic LINE-1 retrotransposition in cortical neurons and non-brain tissues of Rett patients and healthy individuals. PLoS Genet 15, e1008043.

17. Coufal, N.G., Garcia-Perez, J.L., Peng, G.E., Marchetto, M.C.N., Muotri, A.R., Mu, Y., Carson, C.T., Macia, A., Moran, J.V., and Gage, F.H. (2011). Ataxia telangiectasia mutated (ATM) modulates long interspersed element-1 (L1) retrotransposition in human neural stem cells. Proc Natl Acad Sci U S A 108, 20382–20387.

18. Evrony, G.D., Lee, E., Mehta, B.K., Benjamini, Y., Johnson, R.M., Cai, X., Yang, L., Haseley, P., Lehmann, H.S., Park, P.J., et al. (2015). Cell lineage analysis in human brain using endogenous retroelements. Neuron 85, 49–59.

19. Erwin, J.A., Paquola, A.C.M., Singer, T., Gallina, I., Novotny, M., Quayle, C., Bedrosian, T.A., Alves, F.I.A., Butcher, C.R., Herdy, J.R., et al. (2016). L1-associated genomic regions are deleted in somatic cells of the healthy human brain. Nat Neurosci 19, 1583–1591.

20. Zhu, X., Zhou, B., Pattni, R., Gleason, K., Tan, C., Kalinowski, A., Sloan, S., Fiston-Lavier, A.-S., Mariani, J., Petrov, D., et al. (2021). Machine learning reveals bilateral distribution of somatic L1 insertions in human neurons and glia. Nat Neurosci 24, 186–196.

21. Nam, C.H., Youk, J., Kim, J.Y., Lim, J., Park, J.W., Oh, S.A., Lee, H.J., Park, J.W., Won, H., Lee, Y., et al. (2023). Widespread somatic L1 retrotransposition in normal colorectal epithelium. Nature 617, 540–547.

22. Evrony, G.D., Cai, X., Lee, E., Hills, L.B., Elhosary, P.C., Lehmann, H.S., Parker, J.J., Atabay, K.D., Gilmore, E.C., Poduri, A., et al. (2012). Single-neuron sequencing analysis of L1 retrotransposition and somatic mutation in the human brain. Cell 151, 483–496.

23. Burns, K.H. (2017). Transposable elements in cancer. Nat Rev Cancer 17, 415–424.

24. Erwin, J.A., Marchetto, M.C., and Gage, F.H. (2014). Mobile DNA elements in the generation of diversity and complexity in the brain. Nat Rev Neurosci 15, 497–506.

25. Olson, N.D., Wagner, J., Dwarshuis, N., Miga, K.H., Sedlazeck, F.J., Salit, M., and Zook, J.M. (2023). Variant calling and benchmarking in an era of complete human genome sequences. Nat Rev Genet 24, 464–483.

26. Prodanov, T., and Bansal, V. (2020). Sensitive alignment using paralogous sequence variants improves long-read mapping and variant calling in segmental duplications. Nucleic Acids Res. 48, e114.

27. Zhou, W., Emery, S.B., Flasch, D.A., Wang, Y., Kwan, K.Y., Kidd, J.M., Moran, J.V., and Mills, R.E. (2020). Identification and characterization of occult human-specific LINE-1 insertions using long-read sequencing technology. Nucleic Acids Res. 48, 1146–1163.

28. Ebert, P., Audano, P.A., Zhu, Q., Rodriguez-Martin, B., Porubsky, D., Bonder, M.J., Sulovari, A., Ebler, J., Zhou, W., Serra Mari, R., et al. (2021). Haplotype-resolved diverse human genomes and integrated analysis of structural variation. Science 372. 10.1126/science.abf7117.

29. Chu, C., Borges-Monroy, R., Viswanadham, V.V., Lee, S., Li, H., Lee, E.A., and Park, P.J. (2021). Comprehensive identification of transposable element insertions using multiple sequencing technologies. Nat Commun 12, 3836.

30. Chu, C., Zhao, B., Park, P.J., and Lee, E.A. (2020). Identification and Genotyping of Transposable Element Insertions From Genome Sequencing Data. Curr Protoc Hum Genet 107, e102.

31. Tubio, J.M.C., Li, Y., Ju, Y.S., Martincorena, I., Cooke, S.L., Tojo, M., Gundem, G., Pipinikas, C.P., Zamora, J., Raine, K., et al. (2014). Mobile DNA in cancer. Extensive transduction of nonrepetitive DNA mediated by L1 retrotransposition in cancer genomes. Science 345, 1251343.

32. Faulkner, G.J., and Garcia-Perez, J.L. (2017). L1 Mosaicism in Mammals: Extent, Effects, and Evolution. Trends Genet 33, 802–816.

33. Faulkner, G.J., and Billon, V. (2018). L1 retrotransposition in the soma: a field jumping ahead. Mob DNA 9, 22.

34. Chen, J., Cheng, J., Chen, X., Inoue, M., Liu, Y., and Song, C.-X. (2022). Whole-genome long-read TAPS deciphers DNA methylation patterns at base resolution using PacBio SMRT sequencing technology. Nucleic Acids Res 50, e104.

35. Simpson, J.T., Workman, R.E., Zuzarte, P.C., David, M., Dursi, L.J., and Timp, W. (2017). Detecting DNA cytosine methylation using nanopore sequencing. Nat Methods 14, 407–410.

36. Chen, X., Baker, D., Dolzhenko, E., Devaney, J.M., Noya, J., Berlyoung, A.S., Brandon, R., Hruska, K.S., Lochovsky, L., Kruszka, P., et al. (2025). Genome-wide profiling of highly similar paralogous genes using HiFi sequencing. Nat. Commun. 16, 2340.

37. Liao, W.-W., Asri, M., Ebler, J., Doerr, D., Haukness, M., Hickey, G., Lu, S., Lucas, J.K., Monlong, J., Abel, H.J., et al. (2023). A draft human pangenome reference. Nature 617, 312–324.

38. Logsdon, G.A., Ebert, P., Audano, P.A., Loftus, M., Porubsky, D., Ebler, J., Yilmaz, F., Hallast, P., Prodanov, T., Yoo, D., et al. (2025). Complex genetic variation in nearly complete human genomes. Nature. 10.1038/s41586-025-09140-6.

39. Ewing, A.D., Smits, N., Sanchez-Luque, F.J., Faivre, J., Brennan, P.M., Richardson, S.R., Cheetham, S.W., and Faulkner, G.J. (2020). Nanopore sequencing enables comprehensive transposable element epigenomic profiling. Mol. Cell 80, 915–928.e5.

40. Xiao, C., Chen, Z., Chen, W., Padilla, C., Colgan, M., Wu, W., Fang, L.-T., Liu, T., Yang, Y., Schneider, V., et al. (2022). Personalized genome assembly for accurate cancer somatic mutation discovery using tumor-normal paired reference samples. Genome Biol 23, 237.

41. Zhou, W., Mumm, C., Gan, Y., Switzenberg, J.A., Wang, J., De Oliveira, P., Kathuria, K., Losh, S.J., McDonald, T.L., Bessell, B., et al. (2024). A personalized multi-platform assessment of somatic mosaicism in the human frontal cortex. bioRxiv, 2024.12. 18.629274. 10.1101/2024.12.18.629274.

42. Zumalave, S., Santamarina, M., Espasandín, N.P., Garcia-Souto, D., Temes, J., Baker, T.M., Pequeño-Valtierra, A., Otero, I., Rodríguez-Castro, J., Oitabén, A., et al. (2024). Synchronous L1 retrotransposition events promote chromosomal crossover early in human tumorigenesis. bioRxiv. 10.1101/2024.08.27.596794.

43. Baillie, J.K., Barnett, M.W., Upton, K.R., Gerhardt, D.J., Richmond, T.A., De Sapio, F., Brennan, P.M., Rizzu, P., Smith, S., Fell, M., et al. (2011). Somatic retrotransposition alters the genetic landscape of the human brain. Nature 479, 534–537.

44. McKerrow, W., Tang, Z., Steranka, J.P., Payer, L.M., Boeke, J.D., Keefe, D., Fenyö, D., Burns, K.H., and Liu, C. (2020). Human transposon insertion profiling by sequencing (TIPseq) to map LINE-1 insertions in single cells. Philos Trans R Soc Lond B Biol Sci 375, 20190335.

45. McDonald, T.L., Zhou, W., Castro, C.P., Mumm, C., Switzenberg, J.A., Mills, R.E., and Boyle, A.P. (2021). Cas9 targeted enrichment of mobile elements using nanopore sequencing. Nat. Commun. 12, 3586.

46. Coorens, T.H.H., Oh, J.W., Choi, Y.A., Lim, N.S., Zhao, B., Voshall, A., Abyzov, A., Antonacci-Fulton, L., Aparicio, S., Ardlie, K.G., et al. (2025). The somatic mosaicism across human tissues network. 643, 47–59.

47. Jiang, T., Liu, Y., Jiang, Y., Li, J., Gao, Y., Cui, Z., Liu, Y., Liu, B., and Wang, Y. (2020). Long-read-based human genomic structural variation detection with cuteSV. Genome Biol 21, 189.

48. Smolka, M., Paulin, L.F., Grochowski, C.M., Horner, D.W., Mahmoud, M., Behera, S., Kalef-Ezra, E., Gandhi, M., Hong, K., Pehlivan, D., et al. (2024). Detection of mosaic and population-level structural variants with Sniffles2. Nat Biotechnol 42, 1571–1580.

49. Tarailo-Graovac, M., and Chen, N. (2009). Using RepeatMasker to identify repetitive elements in genomic sequences. Curr Protoc Bioinformatics *Chapter 4*, 4.10.1–4.10.14.

50. Li, H. (2025). Finding easy regions for short-read variant calling from pangenome data. arXiv [q-bio.GN].

51. Keskus, A.G., Bryant, A., Ahmad, T., Yoo, B., Aganezov, S., Goretsky, A., Donmez, A., Lansdon, L.A., Rodriguez, I., Park, J., et al. (2025). Severus detects somatic structural variation and complex rearrangements in cancer genomes using long-read sequencing. Nat Biotechnol. 10.1038/s41587-025-02618-8.

52. Brouha, B., Schustak, J., Badge, R.M., Lutz-Prigge, S., Farley, A.H., Moran, J.V., and Kazazian, H.H., Jr (2003). Hot L1s account for the bulk of retrotransposition in the human population. Proc Natl Acad Sci U S A 100, 5280–5285.

53. Schloissnig, S., Pani, S., Ebler, J., Hain, C., Tsapalou, V., Söylev, A., Hüther, P., Ashraf, H., Prodanov, T., Asparuhova, M., et al. (2025). Structural variation in 1,019 diverse humans based on long-read sequencing. Nature 644, 442–452.

54. Gardner, E.J., Lam, V.K., Harris, D.N., Chuang, N.T., Scott, E.C., Pittard, W.S., Mills, R.E., 1000 Genomes Project Consortium, and Devine, S.E. (2017). The Mobile Element Locator Tool (MELT): population-scale mobile element discovery and biology. Genome Res 27, 1916–1929.

55. Rodriguez-Martin, B., Alvarez, E.G., Baez-Ortega, A., Zamora, J., Supek, F., Demeulemeester, J., Santamarina, M., Ju, Y.S., Temes, J., Garcia-Souto, D., et al. (2023). Author Correction: Pan-cancer analysis of whole genomes identifies driver rearrangements promoted by LINE-1 retrotransposition. Nat. Genet. 55, 1080.

56. Sievers, F., Wilm, A., Dineen, D., Gibson, T.J., Karplus, K., Li, W., Lopez, R., McWilliam, H., Remmert, M., Söding, J., et al. (2011). Fast, scalable generation of high-quality protein multiple sequence alignments using Clustal Omega. Mol Syst Biol 7, 539.

57. Ostertag, E.M., and Kazazian, H.H., Jr (2001). Twin priming: a proposed mechanism for the creation of inversions in L1 retrotransposition. Genome Res 11, 2059–2065.

58. Buzdin, A., Gogvadze, E., Kovalskaya, E., Volchkov, P., Ustyugova, S., Illarionova, A., Fushan, A., Vinogradova, T., and Sverdlov, E. (2003). The human genome contains many types of chimeric retrogenes generated through in vivo RNA recombination. Nucleic Acids Res 31, 4385–4390.

59. Garcia-Perez, J.L., Doucet, A.J., Bucheton, A., Moran, J.V., and Gilbert, N. (2007). Distinct mechanisms for trans-mediated mobilization of cellular RNAs by the LINE-1 reverse transcriptase. Genome Res 17, 602–611.

60. Doucet, A.J., Droc, G., Siol, O., Audoux, J., and Gilbert, N. (2015). U6 snRNA Pseudogenes: Markers of Retrotransposition Dynamics in Mammals. Mol Biol Evol 32, 1815–1832.

61. Law, C.-T., and Burns, K.H. (2025). Comparative Genomics Reveals LINE-1 Recombination with Diverse RNAs. bioRxiv. 10.1101/2025.02.02.635956.

62. Chuang, N.T., Gardner, E.J., Terry, D.M., Crabtree, J., Mahurkar, A.A., Rivell, G.L., Hong, C.C., Perry, J.A., and Devine, S.E. (2021). Mutagenesis of human genomes by endogenous mobile elements on a population scale. Genome Res 31, 2225–2235.

63. Aricescu, A.R., Siebold, C., Choudhuri, K., Chang, V.T., Lu, W., Davis, S.J., van der Merwe, P.A., and Jones, E.Y. (2007). Structure of a tyrosine phosphatase adhesive interaction reveals a spacer-clamp mechanism. Science 317, 1217–1220.

64. Liu, W., Liu, Y., Zhang, L., Li, L., Yang, W., Li, J., and He, W. (2025). Nucleic acid spheres for treating capillarisation of liver sinusoidal endothelial cells in liver fibrosis. Nat Commun 16, 4517.

65. Luo, Z., Mao, X., and Cui, W. (2019). Circular RNA expression and circPTPRM promotes proliferation and migration in hepatocellular carcinoma. Med Oncol 36, 86.

66. Koiliari, E., Roussos, P., Pasparakis, E., Lencz, T., Malhotra, A., Siever, L.J., Giakoumaki, S.G., and Bitsios, P. (2014). The CSMD1 genome-wide associated schizophrenia risk variant rs10503253 affects general cognitive ability and executive function in healthy males. Schizophr Res 154, 42–47.

67. Byrne, R.A.J., Nimmo, J., Torvell, M., Carpanini, S.M., Daskoulidou, N., Hughes, T.R., Noble, L.V., Veteleanu, A., Watkins, L.M., Zelek, W.M., et al. (2025). The schizophrenia-associated gene CSMD1 encodes a complement classical pathway inhibitor predominantly expressed by astrocytes and at synapses in mice and humans. Brain Behav Immun 127, 287–302.

68. Athanasiu, L., Giddaluru, S., Fernandes, C., Christoforou, A., Reinvang, I., Lundervold, A.J., Nilsson, L.-G., Kauppi, K., Adolfsson, R., Eriksson, E., et al. (2017). A genetic association study of CSMD1 and CSMD2 with cognitive function. Brain Behav Immun 61, 209–216.

69. Park, J., Cook, D.E., Chang, P.-C., Kolesnikov, A., Brambrink, L., Mier, J.C., Gardner, J., McNulty, B., Sacco, S., Keskus, A., et al. (2024). DeepSomatic: Accurate somatic small variant discovery for multiple sequencing technologies. bioRxivorg. 10.1101/2024.08.16.608331.

70. Antipov, D., Rautiainen, M., Nurk, S., Walenz, B.P., Solar, S.J., Phillippy, A.M., and Koren, S. (2025). Verkko2 integrates proximity-ligation data with long-read De Bruijn graphs for efficient telomere-to-telomere genome assembly, phasing, and scaffolding. Genome Res. 35, 1583–1594.

71. Stanojevic, D., Lin, D., Florez De Sessions, P., and Sikic, M. (2024). Telomere-to-telomere phased genome assembly using error-corrected Simplex nanopore reads. bioRxiv, 2024.05.18.594796. 10.1101/2024.05.18.594796.

72. Li, H., Bloom, J.M., Farjoun, Y., Fleharty, M., Gauthier, L., Neale, B., and MacArthur, D. (2018). A synthetic-diploid benchmark for accurate variant-calling evaluation. Nat. Methods 15, 595–597.

73. Manni, M., Berkeley, M.R., Seppey, M., Simão, F.A., and Zdobnov, E.M. (2021). BUSCO update: Novel and streamlined workflows along with broader and deeper phylogenetic coverage for scoring of eukaryotic, prokaryotic, and viral genomes. Mol. Biol. Evol. 38, 4647–4654.

74. Mikheenko, A., Prjibelski, A., Saveliev, V., Antipov, D., and Gurevich, A. (2018). Versatile genome assembly evaluation with QUAST-LG. Bioinformatics 34, i142–i150.

75. Rhie, A., Walenz, B.P., Koren, S., and Phillippy, A.M. (2020). Merqury: reference-free quality, completeness, and phasing assessment for genome assemblies. Genome Biol. 21, 245.

76. Storer, J., Hubley, R., Rosen, J., Wheeler, T.J., and Smit, A.F. (2021). The Dfam community resource of transposable element families, sequence models, and genome annotations. Mob DNA 12, 2.

77. Fernandes, J.D., Zamudio-Hurtado, A., Clawson, H., Kent, W.J., Haussler, D., Salama, S.R., and Haeussler, M. (2020). The UCSC repeat browser allows discovery and visualization of evolutionary conflict across repeat families. Mob. DNA 11, 13.

78. Shiraishi, Y., Koya, J., Chiba, K., Okada, A., Arai, Y., Saito, Y., Shibata, T., and Kataoka, K. (2023). Precise characterization of somatic complex structural variations from tumor/control paired long-read sequencing data with nanomonsv. Nucleic Acids Res 51, e74.

79. Nam, C.H., Youk, J., Kim, J.Y., Lim, J., Park, J.W., Oh, S.A., Lee, H.J., Park, J.W., Won, H., Lee, Y., et al. (2023). Widespread somatic L1 retrotransposition in normal colorectal epithelium. Nature 617, 540–547.

80. Li, H., Handsaker, B., Wysoker, A., Fennell, T., Ruan, J., Homer, N., Marth, G., Abecasis, G., Durbin, R., and 1000 Genome Project Data Processing Subgroup (2009). The Sequence Alignment/Map format and SAMtools. Bioinformatics 25, 2078–2079.

81. 10.5281/zenodo.15733390.

82. Camacho, C., Coulouris, G., Avagyan, V., Ma, N., Papadopoulos, J., Bealer, K., and Madden, T.L. (2009). BLAST+: architecture and applications. BMC Bioinformatics 10, 421.

83. R Core Team (2024). R: A Language and Environment for Statistical Computing.

84. Cheng, H., Concepcion, G.T., Feng, X., Zhang, H., and Li, H. (2021). Haplotype-resolved de novo assembly using phased assembly graphs with hifiasm. Nat Methods 18, 170–175.

85. Niu, Y., Teng, X., Zhou, H., Shi, Y., Li, Y., Tang, Y., Zhang, P., Luo, H., Kang, Q., Xu, T., et al. (2022). Characterizing mobile element insertions in 5675 genomes. Nucleic Acids Res 50, 2493–2508.

86. Kojima, S., Koyama, S., Ka, M., Saito, Y., Parrish, E.H., Endo, M., Takata, S., Mizukoshi, M., Hikino, K., Takeda, A., et al. (2023). Mobile element variation contributes to population-specific genome diversification, gene regulation and disease risk. Nat Genet 55, 939–951.

87. Borges-Monroy, R., Chu, C., Dias, C., Choi, J., Lee, S., Gao, Y., Shin, T., Park, P.J., Walsh, C.A., and Lee, E.A. (2021). Whole-genome analysis reveals the contribution of non-coding de novo transposon insertions to autism spectrum disorder. Mob DNA 12, 28.

88. Collins, R.L., Brand, H., Karczewski, K.J., Zhao, X., Alföldi, J., Francioli, L.C., Khera, A.V., Lowther, C., Gauthier, L.D., Wang, H., et al. (2020). A structural variation reference for medical and population genetics. Nature 581, 444–451.

89. Robinson, J.T., Thorvaldsdóttir, H., Winckler, W., Guttman, M., Lander, E.S., Getz, G., and Mesirov, J.P. (2011). Integrative genomics viewer. Nat. Biotechnol. 29, 24–26.

90. Kent, W.J. (2002). BLAT--the BLAST-like alignment tool. Genome Res. 12, 656–664.

91. Wang, S., Zhou, W., and Wang, J. (2025). SMaHT-MEI-Benchmarking. (Zenodo). 10.5281/ZENODO.17254344.

