## Supplements for "Multi-platform framework for mapping somatic retrotransposition in human tissues": Supplementary Information.pdf

Figure S1. sMEI benchmarking set pipeline in HapMap mixture
Figure S2. Statistics of sMEI benchmarking set in HapMap mixture
Figure S3. sMEI detection method benchmark in HapMap mixture (L1, tier1)
Figure S4. sMEI detection method benchmark in HapMap mixture (*Alu* and SVA, tier2)
Figure S5. The Recall and precision of integrative strategy in HapMap mixture (Illumina and PacBio)
Figure S6. Systematic analysis of TP and FP compositions in WGS-based sMEI detection methods
Figure S7. The F1, recall and precision of integrative strategy in HapMap mixture (Illumina and ONT)
Figure S8. The F1, recall and precision of integrative strategy using independent short-read and long-read WGS data in HapMap mixture (Four other GCCs)
Figure S9. DSA generation pipeline
Figure S10. Allele Frequencies across benchmarking sets and samples
Figure S11. Performance evaluation across refinement methods and mixture levels
Figure S12. IGV screenshots for two false positive calls at chr11:4,276,716 and chr11:4291927 within segmental duplication regions across GRCh38 and DSA
Figure S13. Source tracing using L1 internal sequence
Figure S14. IGV screenshots for high-confidence L1 insertions in donor tissue samples supported by both short- and long-reads
Figure S15. PCR validation of putative somatic L1 insertions in donor tissue samples
Figure S16. IGV screenshots for high-confidence L1 insertions captured by TEnCATS

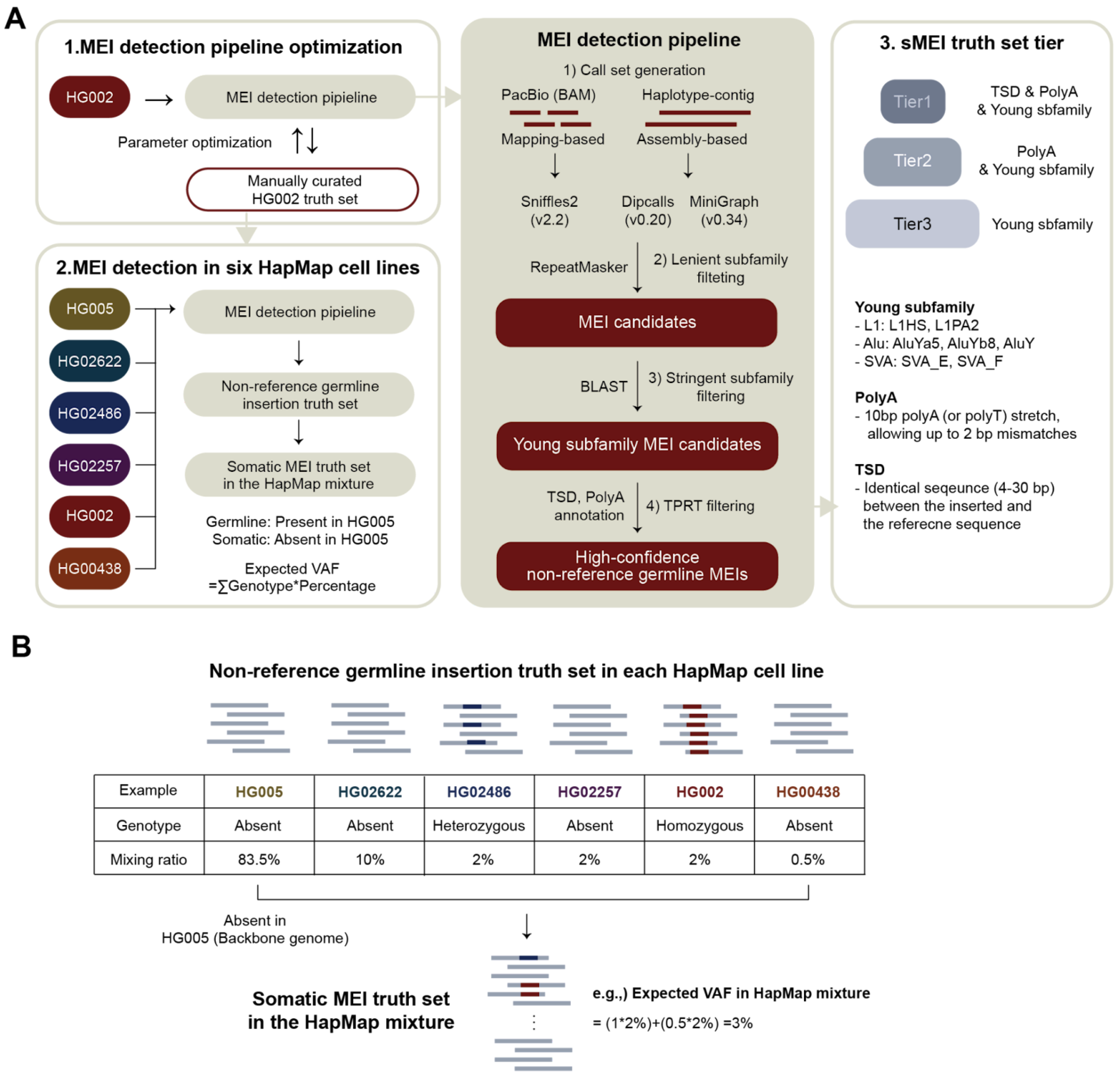


**Figure S1. sMEI benchmarking set pipeline in HapMap mixture A**, A schematic overview of the sMEI benchmarking set generation pipeline. Germline MEIs in each cell line were first detected using long-read WGS (PacBio) data and haplotype contigs, followed by subfamily annotation using RepeatMasker and BLAST. Target-primed reverse transcription (TPRT) features—including target site duplications (TSDs) and poly(A) tails—were then applied to generate high-confidence non-reference germline MEIs in each cell line. MEIs absent in the backbone genome (HG005) but present in any of the other five cell lines were considered as somatic MEIs in the HapMap mixtures. **B**, Illustration of mixture VAF calculation in the HapMap mixtures based on mixing ratios and genotypes. For example, a heterozygous insertion in HG02486 (2%) and homozygous insertion in HG002 (2%) results in a mixture VAF of 3%.

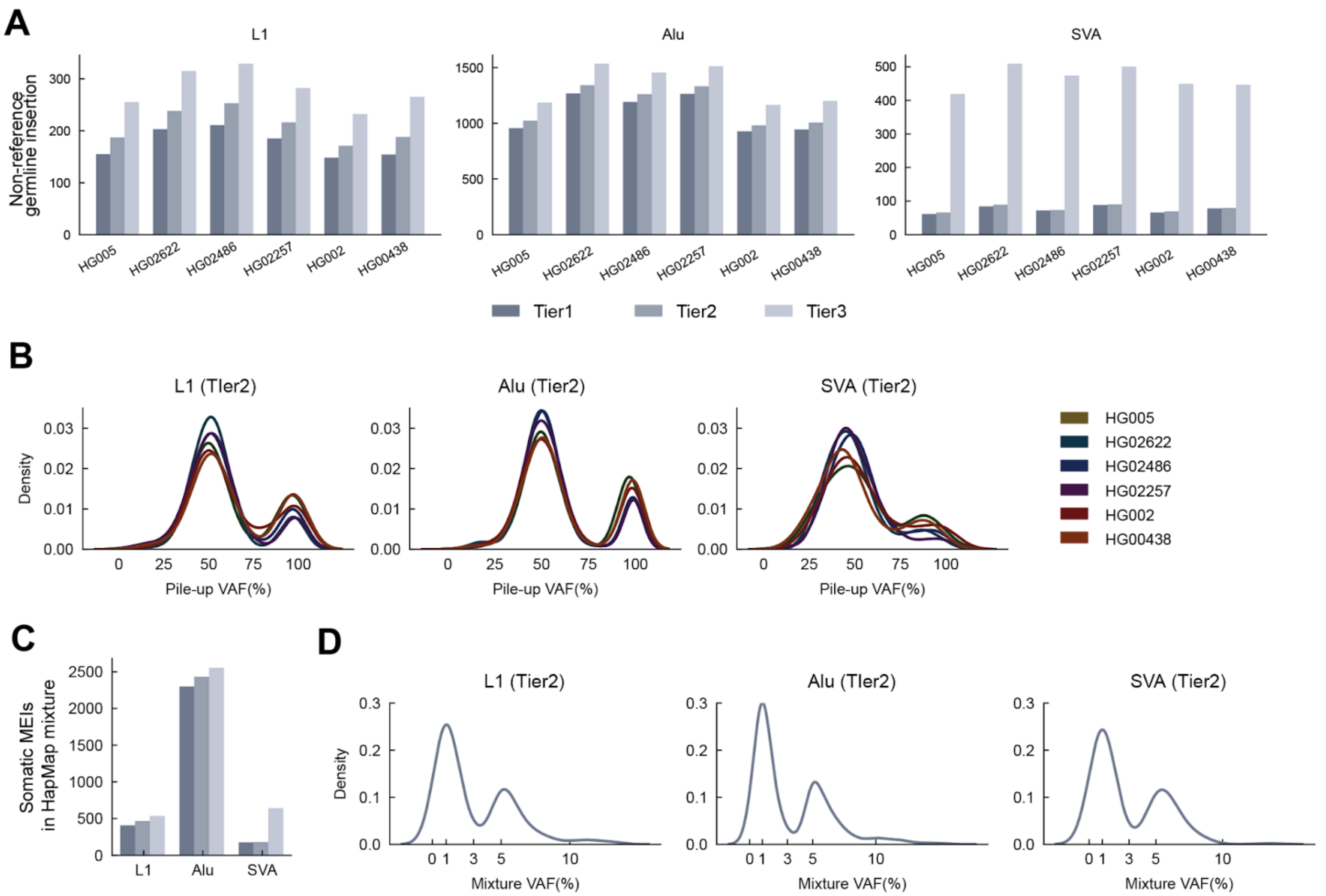


**Figure S2. Statistics of sMEI benchmarking set in HapMap mixture A**, Number of non-reference germline MEIs detected in each of the six HapMap cell lines (HG005, HG02622, HG02486, HG02257, HG002, and HG00438), stratified by element type (L1, *Alu*, and SVA) and benchmarking set tier. The numbers are also provided in Table S4. **B**, Distribution of pile-up VAFs (%) of non-reference germline MEIs for L1, *Alu*, and SVA across the six cell lines in the tier 2 benchmarking set. Each line represents a different cell line. **C**, Number of sMEIs in the HapMap mixtures stratified by element type and benchmarking set tier. **D**, Density distribution of mixture VAFs (%) of L1, *Alu*, and SVA in the tier 2 benchmarking set.

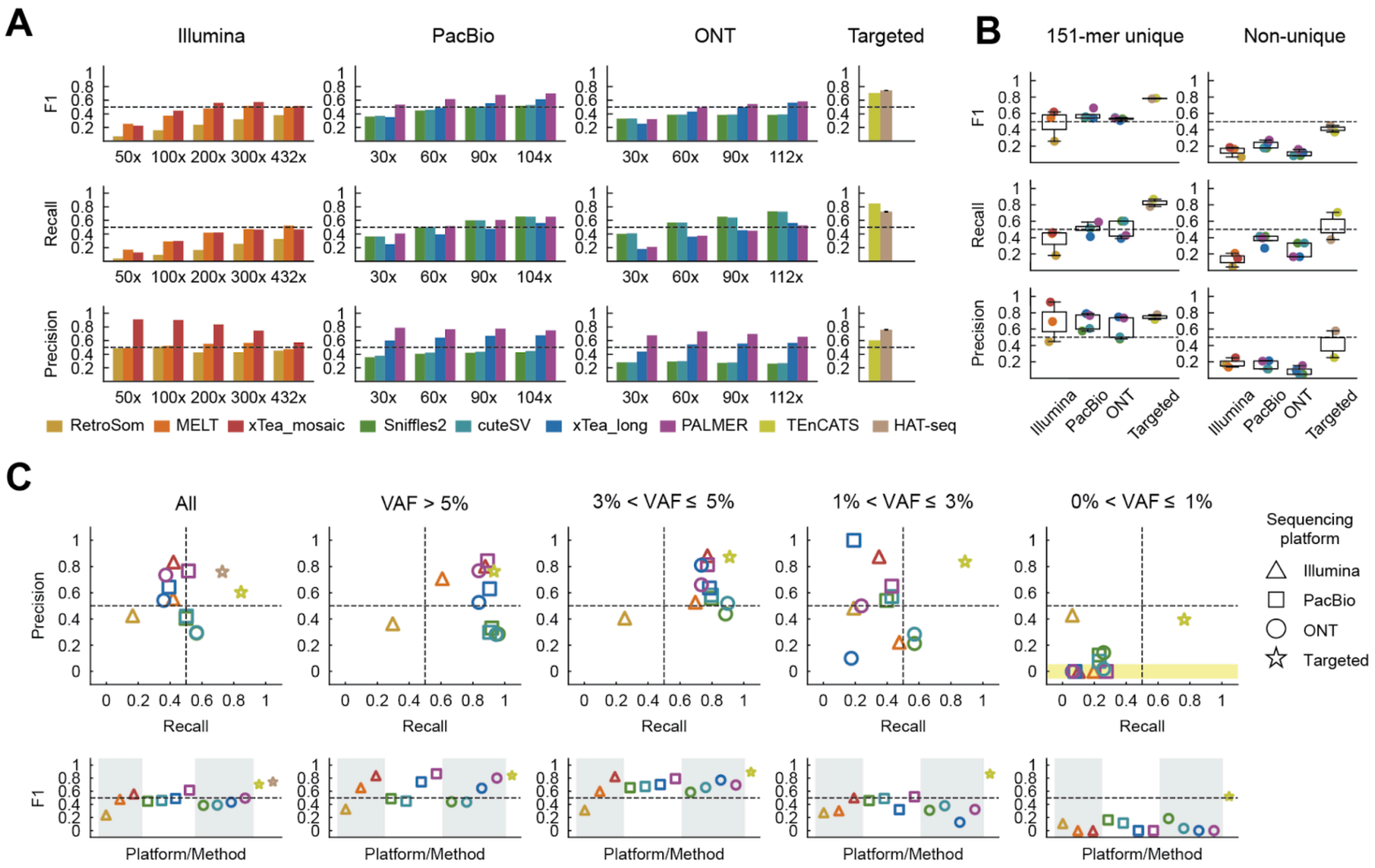


**Figure S3. sMEI detection method benchmark in HapMap mixture (L1, tier1) A,** F1 scores, recall, and precision across different sequencing platforms (Illumina, PacBio, ONT, and MEI-targeted sequencing) and sequencing depth. Methods are distinguished by color. In b–d, 200x and 60x callsets were used for short-read (Illumina) and long-read (PacBio and ONT) data, respectively. The performance in HAT-seq is the average of four experimental replicates. **B,** F1 scores, recall, and precision across genomic regions, including 151-mer unique vs. non-unique regions. Methods are distinguished by color. **C,** F1 scores, recall, and precision in each VAF bin. Platforms are indicated by shape and methods by color. The dots located in the gray area in the recall-precision scatter plot indicate that the corresponding method does not report the calls in the corresponding VAF bin. For HAT-seq, only recall is shown because caller-level VAFs are not available for precision calculation.

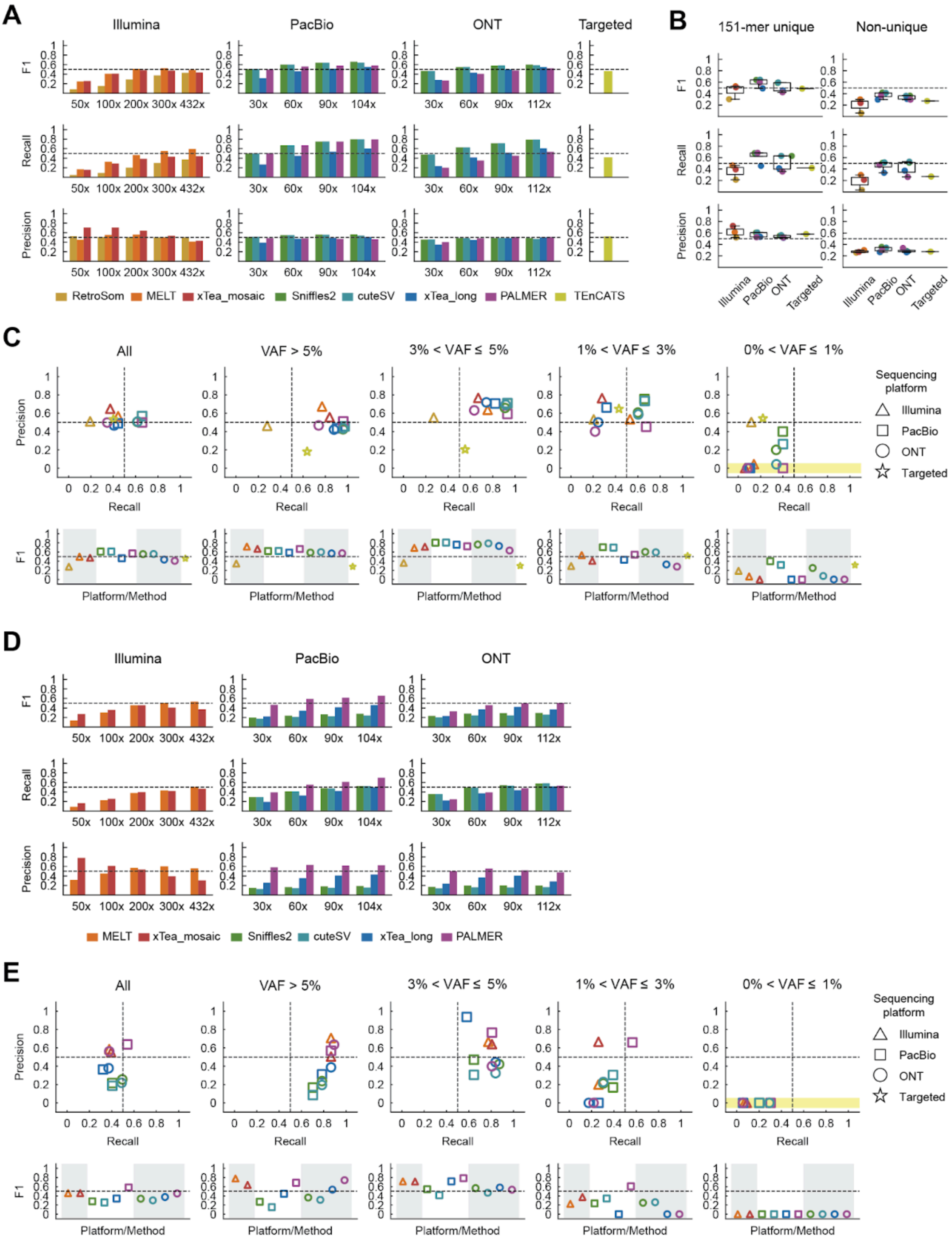
A
Illumina
PacBio
ONT
Targeted
F1
Recall
Precision
50x 100x 200x 300x 432x
30x 60x 90x 104x
30x 60x 90x 112x
RetroSom MELT xTea_mosaic Sniffles2 cuteSV xTea_long PALMER TEnCATS
B
151-mer unique
Non-unique
Illumina PacBio ONT Targeted
C
All
VAF > 5%
3% < VAF ≤ 5%
1% < VAF ≤ 3%
0% < VAF ≤ 1%
Sequencing platform
Illumina
PacBio
ONT
Targeted
Platform/Method
D
MELT xTea_mosaic Sniffles2 cuteSV xTea_long PALMER
E

1322

**Figure S4. sMEI detection method benchmark in HapMap mixture (*Alu* and SVA, tier2) A** and **D,** F1 scores, recall, and precision across different sequencing platforms (Illumina, PacBio, ONT, and MEI-targeted sequencing) and sequencing depth in *Alu* and SVA, respectively. Methods are distinguished by color. In b–d, 200x and 60x callsets were used for short-read (Illumina) and long-read (PacBio and ONT) data, respectively. **B,** F1 scores, recall, and precision across genomic regions, including 151-mer unique vs. non-unique regions. Methods are distinguished by color. **C** and **D,** F1 scores, recall, and precision in each VAF bin in *Alu* and SVA, respectively. Platforms are indicated by shape and methods by color. The dots located in the yellow area in the recall-precision scatter plot indicate that the corresponding method does not report the calls in the corresponding VAF bin.

In the short-read data for *Alu*, the overall performance pattern for *Alu* was similar to that observed for L1 (**Figure S4A**). We observed that overall performance (F1 score) plateaued at ~200x (xTea_mosaic and MELT), marking it as the recommended depth that balances cost and performance. The performance peaked at 300x, followed by a modest decline beyond this depth such that the difference in performance between 200x and 300x was subtle (<0.02 in F1 score). However, in the long-read data, such F1 scores were achievable at much lower ~30x compared to that they were achievable at ~60x in L1. For instance, MELT showed the best performance (F1 score=0.50) at 200x Illumina among short-read WGS-based methods, a level matched by three long-read WGS-based callers using 30x PacBio; Sniffles2, cuteSV (F1 score=0.51) and PALMER (F1 score=0.50). The long-read (TEnCATS) MEI-targeted sequencing methods performed on par with 200x Illumina and 30x PacBio. It is notable that long-read WGS-based methods using larger than 30x data performed slightly better than TEnCATS, compared to that it showed higher performance than any WGS-based methods in L1.

Like L1, 89.7% and 10.3% of sMEIs fell into the 151-unique and non-unique regions, respectively. In 200x Illumina and 60x PacBio/ONT (as the representative sequencing depth, respectively), the recall in the non-unique region (18.1% on average) in *Alu* was higher than that of L1 (10.8% on average) and long-read WGS-based methods substantially maintained higher recall (2.56-fold in PacBio and 2.33-fold in ONT) compared to the performance from short reads (**Figures S4B**). In addition, when the VAFs reach extremely low (≤1%), long-read WGS-based methods and MEI-targeted sequencing achieved 2.5-fold and 2.0-fold higher recall than short-read WGS-based methods on average, respectively (**Figures S4C**).

In SVA, the performance of 200x Illumina (F1 score=0.46 in xTea_mosaic and MELT) were achievable by PALMER using ~30x PacBio (F1 score=0.47) and ~60x ONT (F1 score=0.45) (**Figure S4D**). In 200x Illumina and 60x PacBio/ONT (as the representative sequencing depth, respectively), when the VAFs reach extremely low (≤1%), long-read WGS-based methods achieved 2.0-fold higher recall than short-read WGS-based methods on average (**Figures S4E**). Since less than 30 sMEIs fell into the non-unique region, we did not include the performance evaluation based on genomic stratification in SVA.

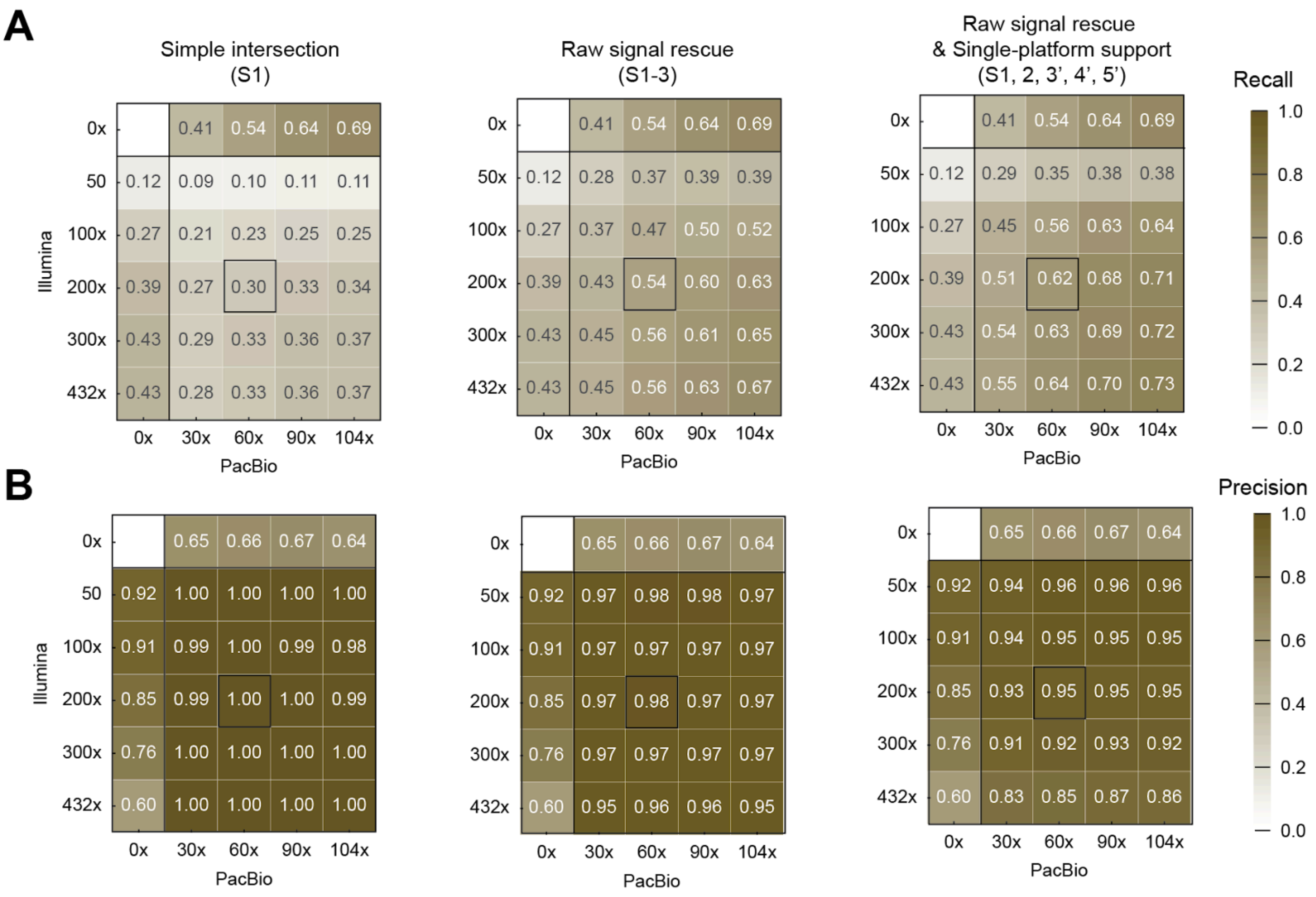


**Figure S5. The recall and precision of integrative strategy (Illumina and PacBio) in HapMap mixture A,** Recall and **B,** precision of integrated callsets across different Illumina and PacBio coverage combinations.

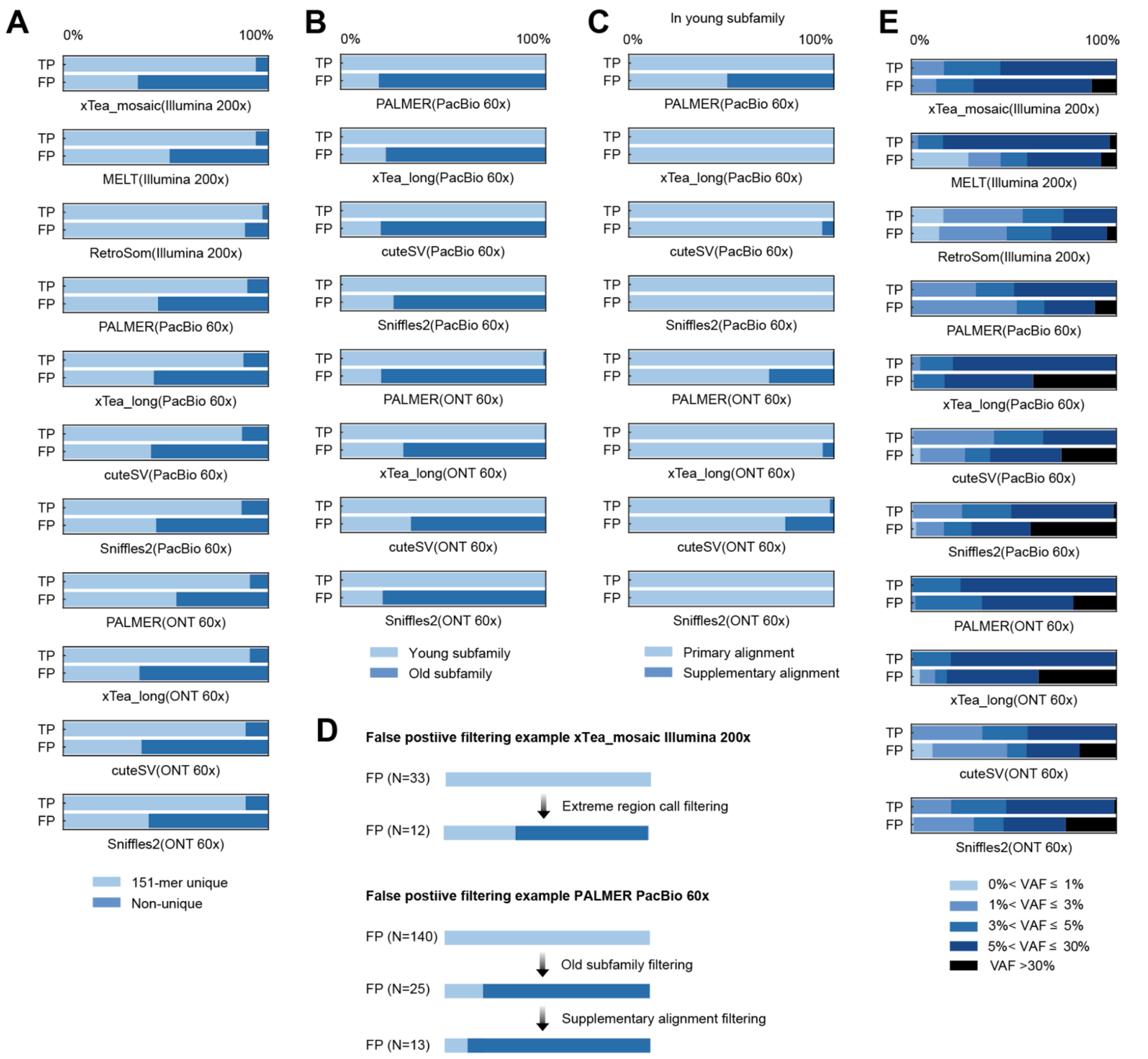


**Figure S6. Systematic analysis of TP and FP compositions in WGS-based sMEI detection methods (Illumina 200x, PacBio and ONT 60x) A,** Percentages (%) of TP and FP belonged to each genomic stratifications (Easy, difficult and extreme region) **B,** Percentages (%) of TP and FP by L1 subfamily in long-read based methods. The young subfamily includes L1HS and L1PA2. **C,** Percentage (%) of TP and FP by supporting reads alignment (primary or supplementary alignment) among the calls of the young subfamily in the long-read based methods. **D,** FP filtering strategy example in xTea_mosaic and PALMER **E**, Percentages (%) of TP and FP by caller VAF levels

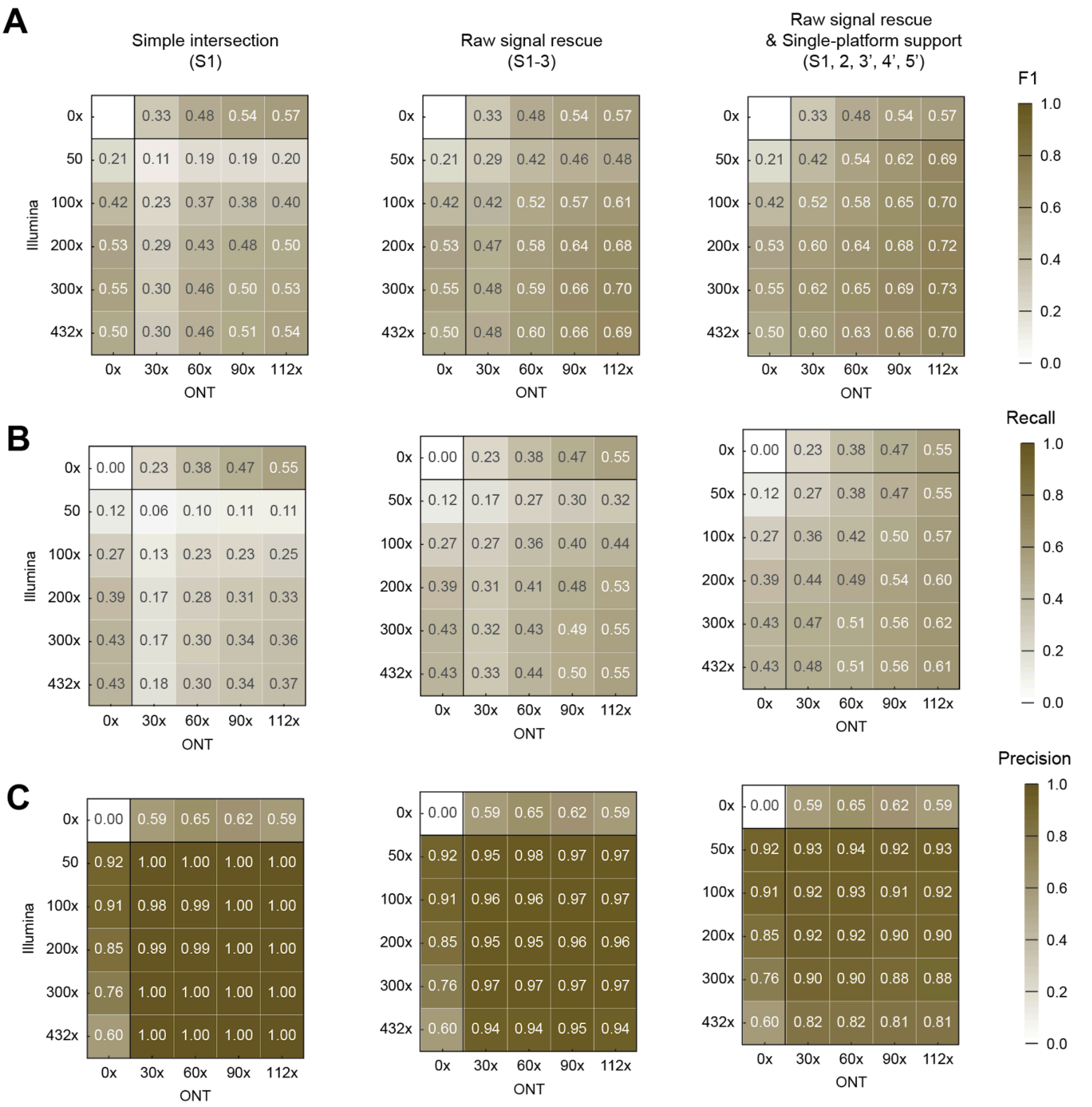


**Figure S7. The F1, recall and precision of integrative strategy (Illumina and ONT) in HapMap mixture A,** F1, **B,** recall and **C,** precision of integrated callsets across different Illumina and ONT coverage combinations.

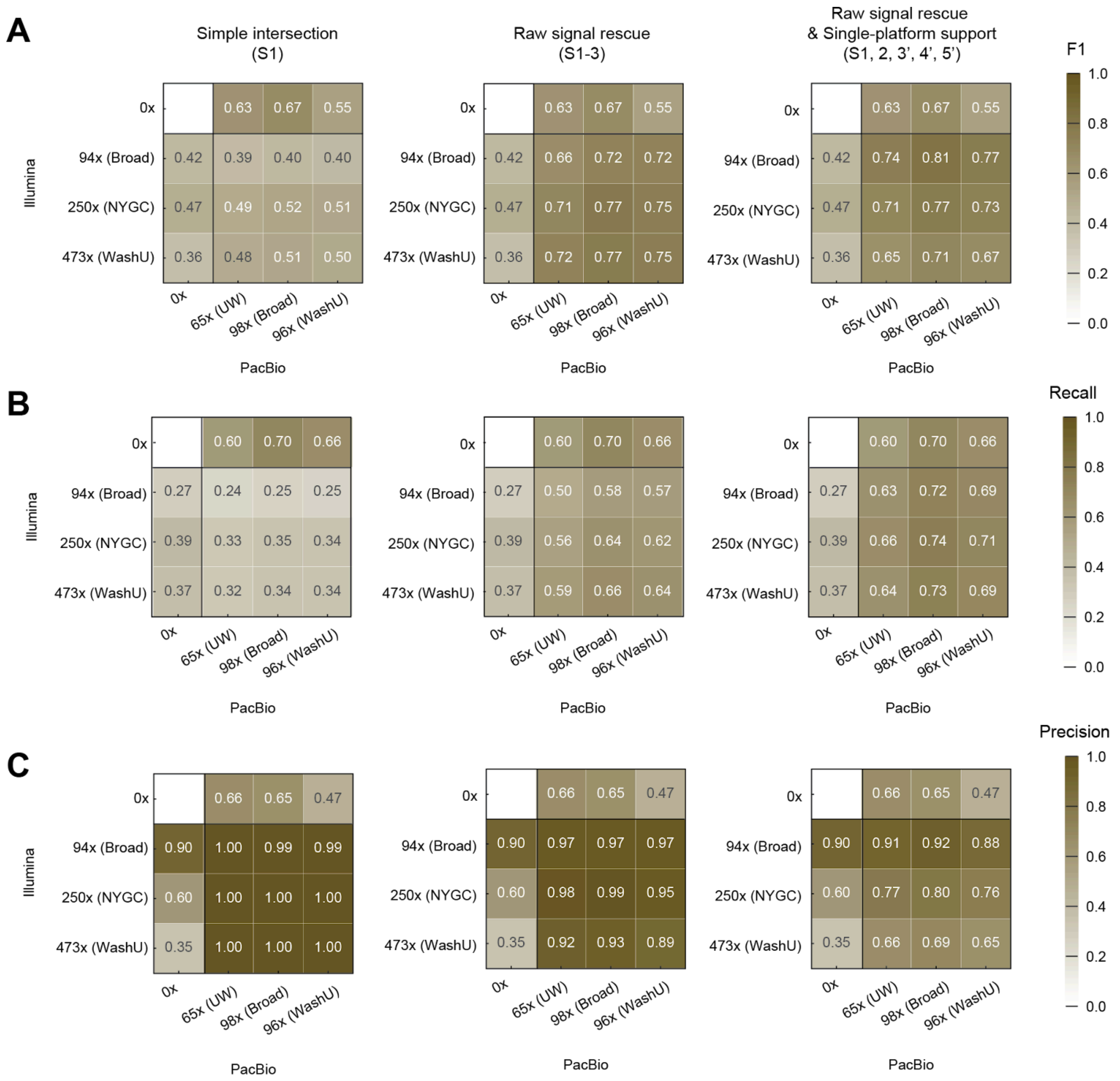


**Figure S8. The F1, recall and precision of integrative strategy using independent short-read and long-read WGS data in HapMap mixture (Four other GCCs) A,** F1, **B,** recall and **C,** precision of integrated callsets across different Illumina and PacBio coverage combinations using independent WGS data from four other GCCs; 94x Illumina and 98x PacBio data from Broad GCC, 250x Illumina from NYGC GCC, 65x PacBio from UW GCC, 473x Illumina and 96x PacBio from WashU GCC.

A

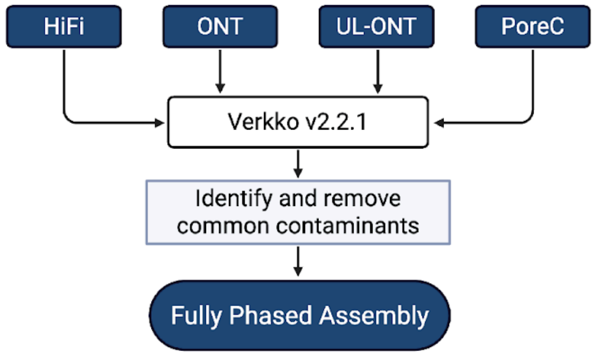


B

| | PacBio Hi-FI | Nanopore ONT | Nanopore UL-ONT | Nanopore Pore-C |
|---|---|---|---|---|
| **Sequence Coverage** | 68x | 30x | 49x | 32x |

C

| | **BL2009** | | **Mean of 65 samples from HGSVC** | | **SMHT012** | |
|---|---|---|---|---|---|---|
| | Verkko v2.2.1 | | Verkko v1.4.1 | | Verkko v2.2.1 | |
| | Hap1 | Hap2 | Hap1 | Hap2 | Hap1 | Hap2 |
| **Total Bases (Gbp)** | 2.997 | 3.024 | 2.985 | 3.005 | 2.929 | 3.022 |
| **N50 (Mbp)** | 145.43 | 136.17 | 133.10 | 137.67 | 135.44 | 140.00 |
| **Number of contigs** | 74 | 102 | 88.4 | 77.4 | 85 | 50 |

D

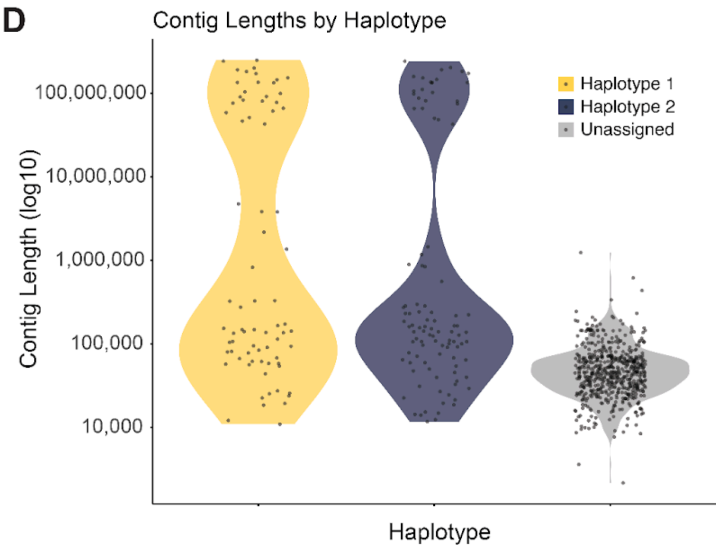


**Figure S9. DSA generation and quality control A,** Donor-specific assembly (DSA) generated using PacBio-HIFI, Nanopore ONT, UL-ONT, and Pore-C sequencing data from the normal cell line (BL2009) in the CASTLE panel project (SRA PRJNA1086849)[69]. These data were collected to generate a DSA without downsampling for this normal-tissue pair using Verkko[70] v2.2.1, the resulting DSA then went through a decontamination process through FCS v0.5.5. **B,** Sequencing coverage of datasets used to construct DSA. **C,** Haplotype-resolved assembly statistics between H2009, SMHT012 and the mean of 65 samples form Human Genome Structural Variation Consortium (HGSVC)[38]. **D,** Per-contig length distributions for DSA. Violin plots ($log_{10}$ scale) show lengths for Haplotype 1, Haplotype 2, and unassigned contigs; points are individual contigs. Both haplotypes show a dominant mode near ~100 Mb, whereas unassigned contigs are shorter.

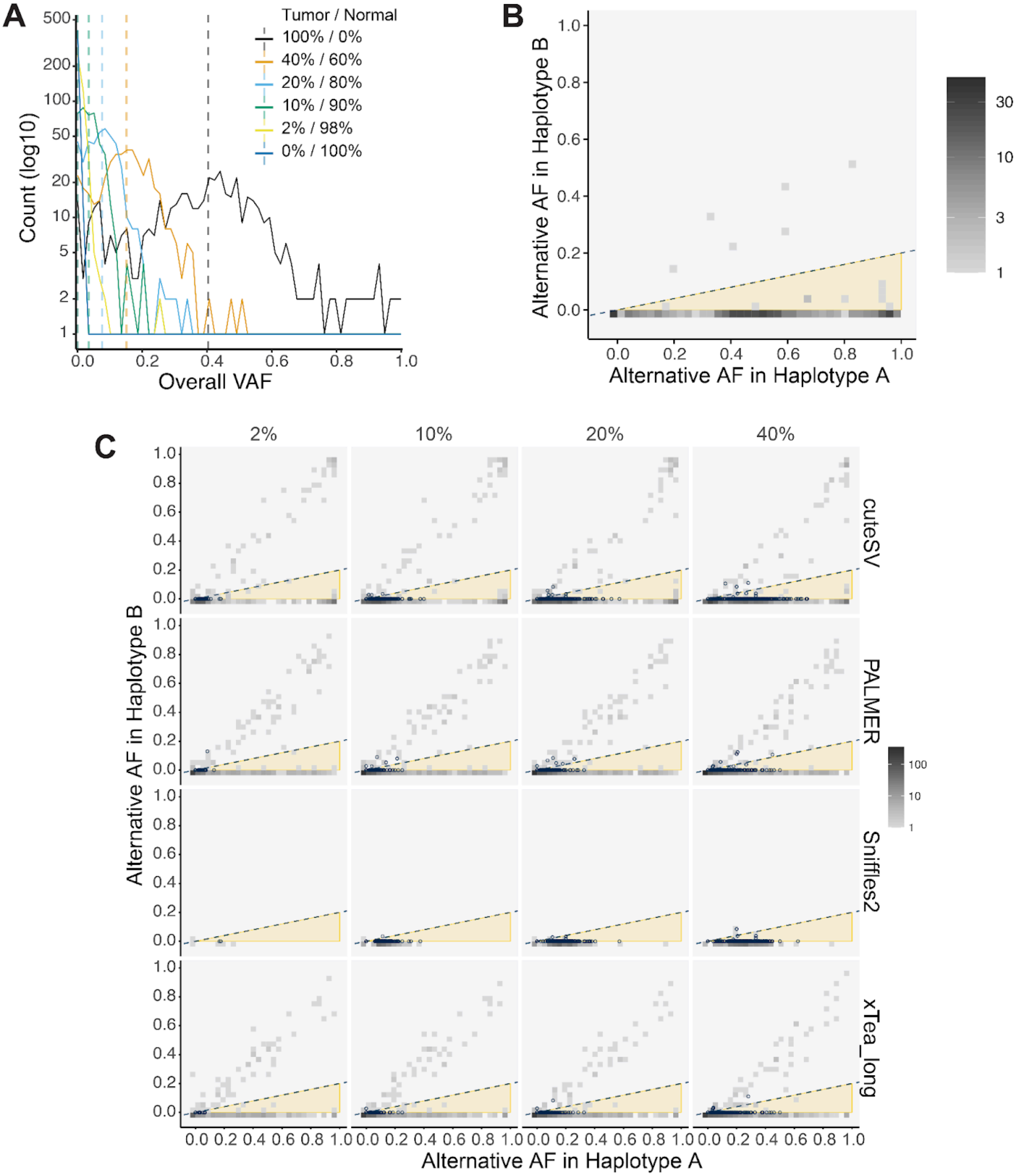


**Figure S10. Allele Frequencies across benchmarking sets and samples A,** Variant allele frequency (VAF) distributions for calls in the cancer benchmarking set across Normal (BL2009), Tumor (H2009), and four mixture (2%, 10%, 20% and 40%) samples using PacBio. **B,** Alternate allele frequencies in Haplotype A vs. Haplotype B. With Haplotype A defined as the haplotype with higher VAF, and Haplotype B defined as the haplotype with lower VAF. We observe 435/441 = 0.986 recall rate with the empirical cutoff y=0.2x. **C,** Alternate VAF in Haplotype A vs. Haplotype B across detection methods and mixture levels.

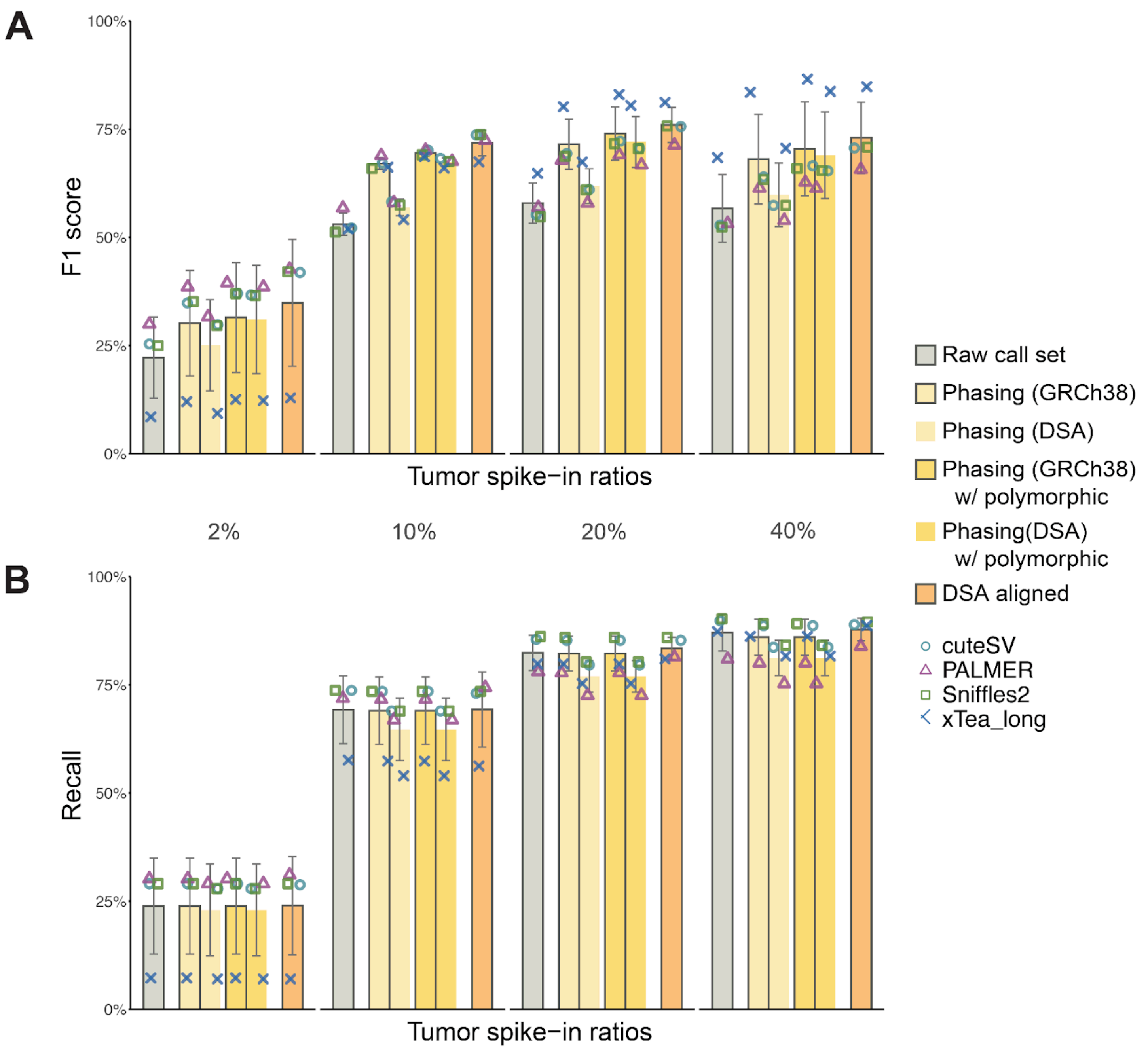


**Figure S11. Performance evaluation across refinement methods and mixture levels A,** F1 score and **B**, recall of sMEI calls across four tumor spike-in mixtures (2%, 10%, 20% and 40%). Individual values from each callers are indicated by colored points.

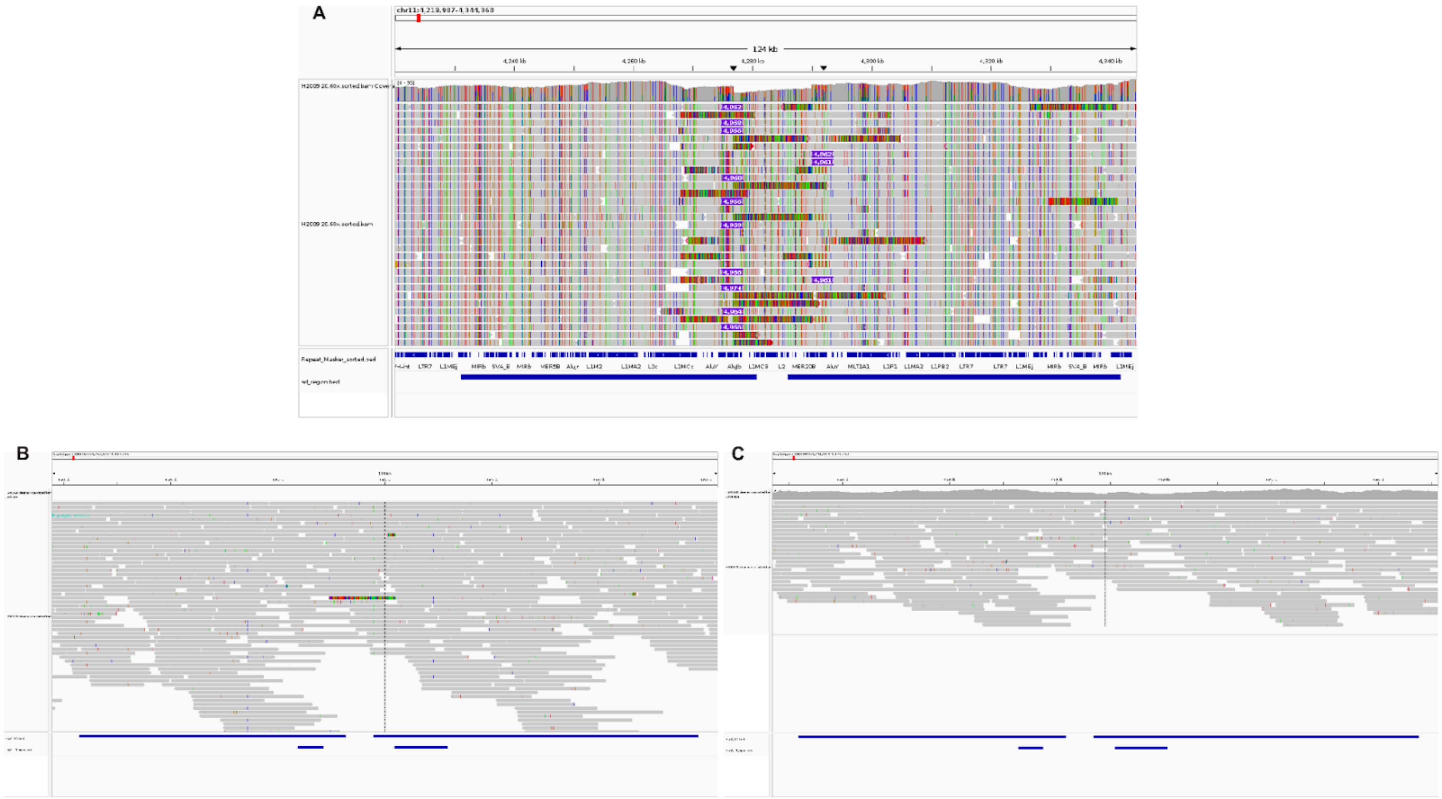


**Figure S12. IGV screenshots for two false positive calls at chr11:4,276,716 and chr11:4291927 within segmental duplication regions across GRCh38 and DSA.**

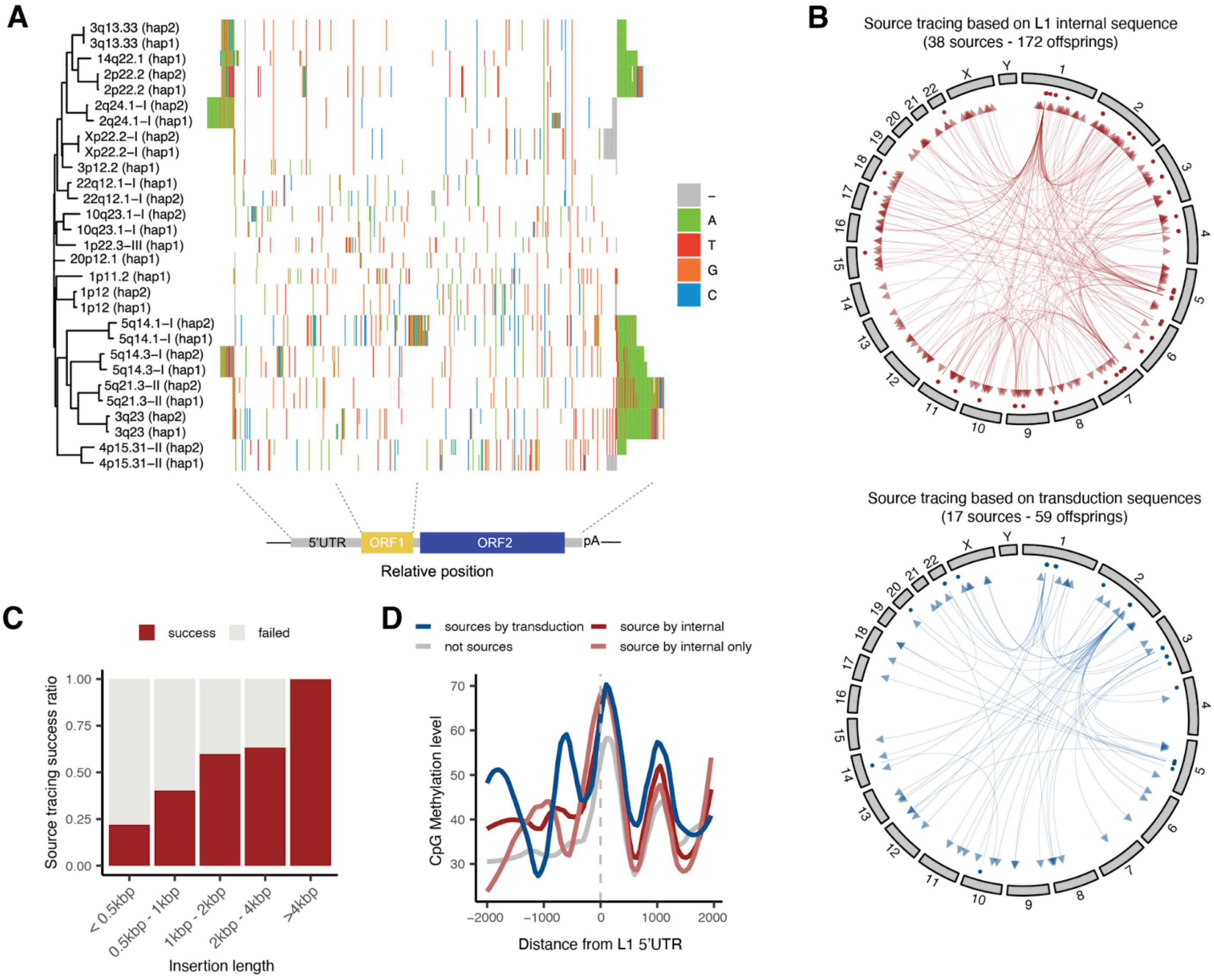


**Figure S13. Source tracing using L1 internal sequence A,** Haplotype-resolved sequence variation in 17 source L1HS identified via transduction sequence. The L1 sources are clustered based on sequence similarity calculated with ClustalO. For each nucleotide position, the most frequent base is colored white, while the other bases and gaps are colored as follows: A (green), T (red), G (orange), C (blue), and gap (gray). **B**, The success rate of source tracing using insertion length (left) and the internal L1 length (right). **C**, DNA methylation profiles at the L1HS 5'UTR for different source types in the BL2009 matched-normal cell line.

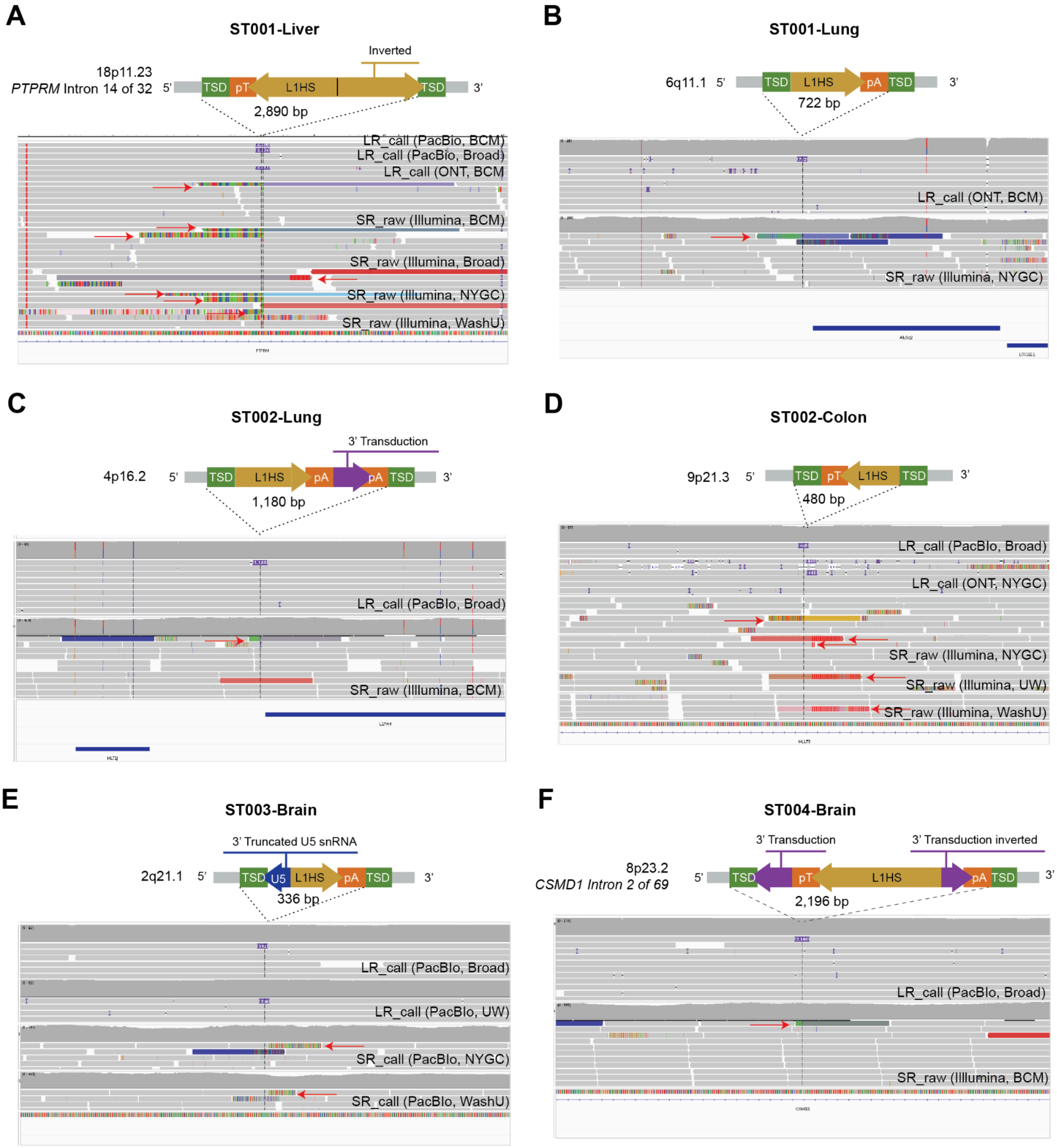


**Figure S14. IGV screenshot for high-confidence L1 insertions in donor tissue samples supported by both short- and long-read**

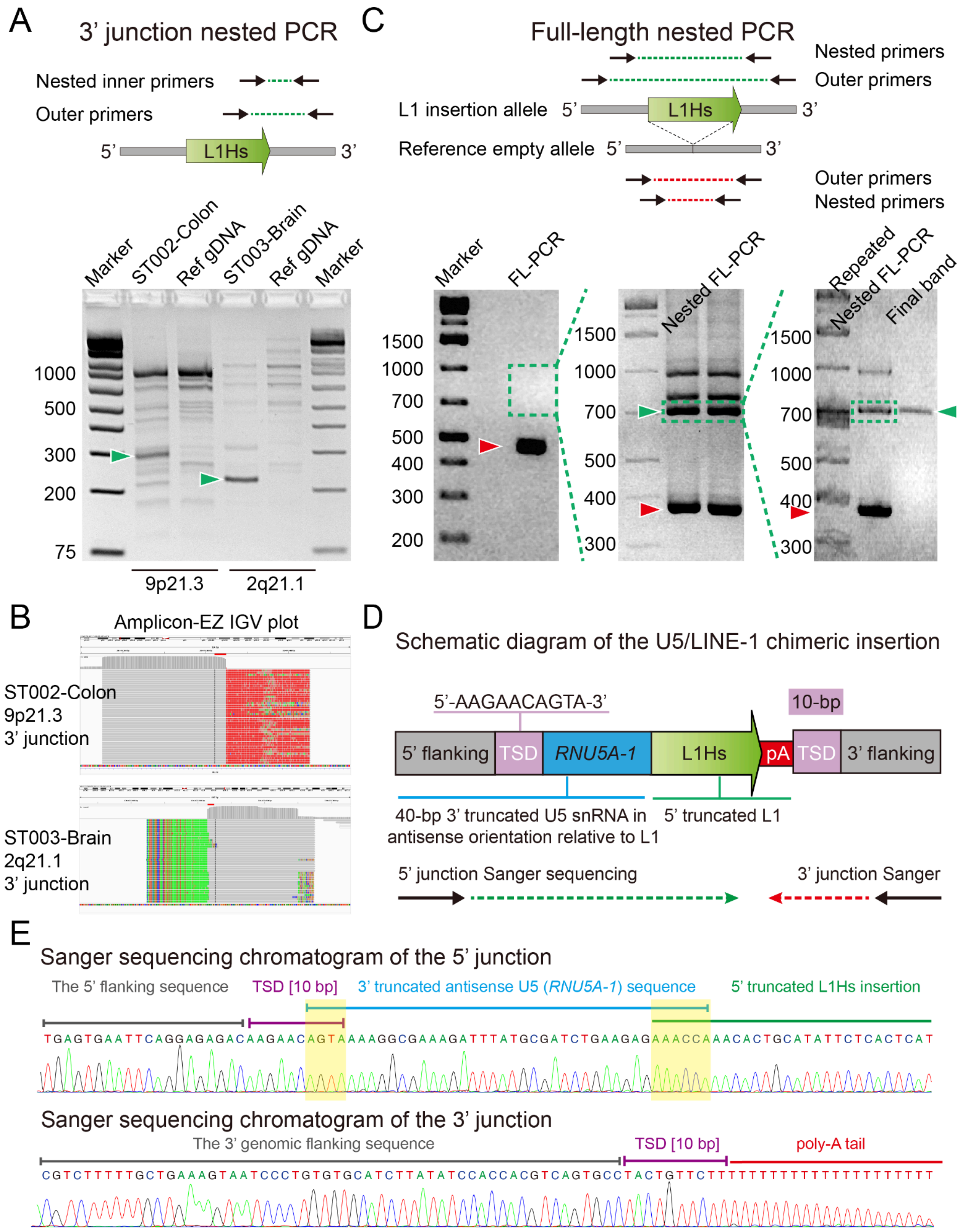
A
3’ junction nested PCR
Nested inner primers
Outer primers
5’
L1Hs
3’
Marker
ST002-Colon
Ref gDNA
ST003-Brain
Ref gDNA
Marker
1000
500
300
200
75
9p21.3
2q21.1
C
Full-length nested PCR
Nested primers
Outer primers
L1 insertion allele 5’
L1Hs
3’
Reference empty allele 5’
3’
Outer primers
Nested primers
Marker
FL-PCR
Nested FL-PCR
Repeated Nested FL-PCR
Final band
1500
1000
700
500
400
300
200
B
Amplicon-EZ IGV plot
ST002-Colon
9p21.3
3’ junction
ST003-Brain
2q21.1
3’ junction
D
Schematic diagram of the U5/LINE-1 chimeric insertion
5’-AAGAACAGTA-3’
10-bp
5’ flanking
TSD
RNU5A-1
L1Hs
pA
TSD
3’ flanking
40-bp 3’ truncated U5 snRNA in antisense orientation relative to L1
5’ truncated L1
5’ junction Sanger sequencing
3’ junction Sanger
E
Sanger sequencing chromatogram of the 5’ junction
The 5’ flanking sequence
TSD [10 bp]
3’ truncated antisense U5 (RNU5A-1) sequence
5’ truncated L1Hs insertion
TGAGTGAATTCAGGAGAGACAAGAACAGTAAAAGGCGAAAGATTTATGCGATCTGAAGAGAAACCAAACACTGCATATTCTCACTCAT
Sanger sequencing chromatogram of the 3’ junction
The 3’ genomic flanking sequence
TSD [10 bp]
poly-A tail
CGTCTTTTTGCTGAAAGTAATCCCTGTGTGCATCTTATATCCACCACGTCAGTGCCTACTGTTCTTTTTTTTTTTTTTTTTTTTTTTTT

**Figure S15. PCR validation of putative somatic L1 insertions in donor tissue samples A**, Schematic of the 3' junction nested PCR strategy (top) and a representative agarose gel (bottom) showing amplification of the 3' junction for two validated somatic L1 insertions, ST002-Colon (9p21.3) and ST003-Brain (2q21.1) tissues. Green arrowheads indicate the target amplicons. Reference genomic DNA (Ref gDNA) was used as a negative control. **B**, Integrative Genomics Viewer (IGV) screenshots showing the alignment of 3' junction sequencing reads (Amplicon-EZ data from Azenta, Inc.) for ST002-Colon (9p21.3) and ST003-Brain (2q21.1) insertion, confirming the somatic L1 integration sites. **C**, Schematic of the full-length nested PCR strategy (top) and representative agarose gels (bottom) for the ST003-Brain (2q21.1) insertion. Due to low mosaicism, the full-length L1 insertion band was not visible after the initial PCR (FL-PCR). A region of the gel at the expected size was excised (dashed blue box) for subsequent DNA extraction and re-amplification. This repeated nested PCR approach successfully isolated a clean band corresponding to the L1 insertion allele (green arrowheads). Red arrowheads indicate the band from the reference empty allele. **D**, Schematic diagram of the U5/L1 chimeric insertion identified in sample ST003-Brain (2q21.1) insertion. The insertion consists of a 40-bp, 3'-truncated U5 small nuclear RNA (snRNA, RNU5A-1) in an antisense orientation relative to a 5'-truncated L1Hs element. The chimeric insertion is flanked by a 10-bp target site duplication (TSD). The locations of primers used for Sanger sequencing are indicated at the bottom. **E**, Sanger chromatograms confirmed the sequence and precise structure of the 5' and 3' junctions of the ST003 brain insertion. The sequences corresponding to the 5' flanking region, 10-bp TSD, truncated U5 sequence, L1Hs element, and 3' poly-A tail are shown. Light yellow rectangles highlight the micro-homology regions between U5 and its flanking genomic and L1Hs sequences.

### Candidate 1: ST001-Liver 18p11.23

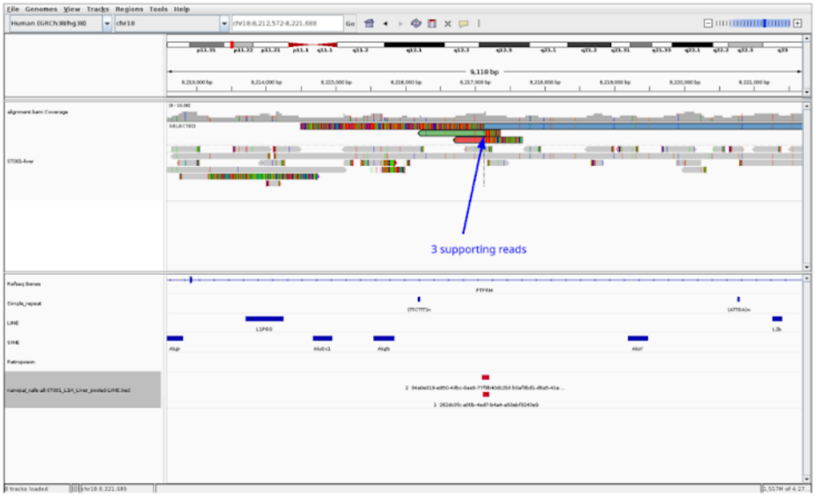


### Candidate 2: ST002-Colon 9p21.3

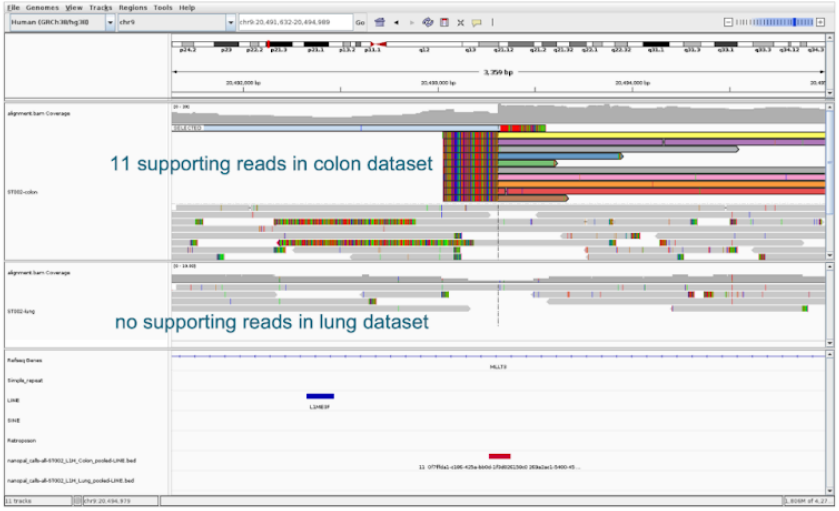


### Candidate 3: ST003-Brain 2q21.1

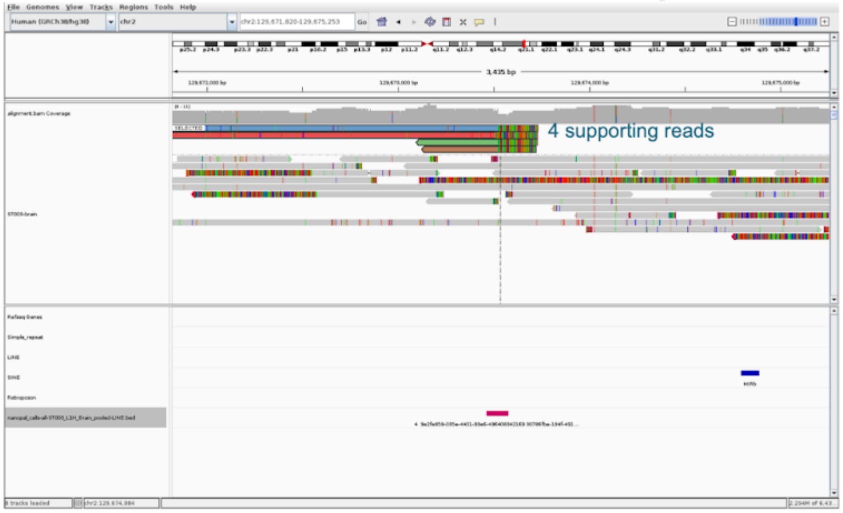


**Figure S16. IGV screenshots for high-confidence L1 insertions captured by TEnCATS.**
